# Reengineering the antigen optimization process for superior neoantigen vaccine design

**DOI:** 10.1101/2025.08.03.668363

**Authors:** Guanqiao Zhang, Yaqi Fu, Damiano Buratto, Kevin C. Chan, Hanyao Wang, Yuanling Huang, Liquan Huang, Ruhong Zhou

**Affiliations:** College of Physics, Institute of Quantitative Biology, and College of Life Sciences, Zhejiang University, Hangzhou 310027, China; Zhejiang Key Laboratory of Cell and Molecular Intelligent Design and Development, Zhejiang University, Hangzhou 310058, China; The First Affiliated Hospital, School of Medicine, Zhejiang University, Hangzhou 310058, China; Department of Chemistry, Columbia University, New York, NY 10027, USA; Department of Biosciences and Bioinformatics, School of Science, Xi’an Jiaotong-Liverpool University, Suzhou, China; Institute WUT-AMU, Wuhan University of Technology, Wuhan 430070, China

**Author notes:** These authors contribute equally to the work (Guanqiao Zhang, Yaqi Fu).

## Abstract

Identifying effective neoantigen sequences is essential for enhancing anti-tumor immunity. However, the vast sequence space (>10^9^ possible peptides) and limited accuracy of existing immunogenicity predictors hinder efficient vaccine design for patient-specific human leukocyte antigens (HLAs). We present AlphaVacc, a deep reinforcement learning framework that integrates Monte Carlo Tree Search with a Transformer-based network to optimize antigenic peptides. AlphaVacc outperforms previous generative models in binding-affinity prediction. Experimental validation of 12 AlphaVacc-generated variants of the BING-4 peptide confirmed that 11 showed increased HLA-A*02:01 binding and 7 elicited significant T cell responses. Further testing of 16 single-mutation peptides confirmed computational predictions for 15 candidates, exhibiting a remarkable success rate. AlphaVacc thus provides a powerful tool for designing neoantigen-based cancer vaccines and may accelerate personalized immunotherapies.

## Introduction

Cancer remains a major global public health challenge. In 2022, nearly 20 million new cancer cases were diagnosed and approximately 9.7 million cancer-related deaths occurred^1^. Nearly one in five individuals will face a cancer diagnosis during their lifetime^1,2^, underscoring the urgent need for effective treatment strategies. Cancer immunotherapy, which empowers patients’ immune systems to recognize and eliminate tumors, has emerged as a powerful approach^3–6^. Several immunotherapies, including FDA-approved immune checkpoint inhibitors (ICI), have demonstrated clinical benefit, although response rates remain limited (∼12.5% overall)^7^. T-cell receptor (TCR)-engineered T cell therapy (TCR-T), such as TECELRA, is currently approved only for synovial sarcoma, a rare cancer type^8^. These limitations highlight the need for novel immunotherapeutic strategies^9–11^.

Neoantigen-based cancer vaccines, using peptides derived from tumor-specific mutations, were shown to be safe and feasible in clinical trials^12–14^. These vaccines rely on peptides (8-10 amino acids) that bind human leukocyte antigen (HLA) class I molecules, forming peptide–HLA (pHLA) complexes recognized by CD8□ T cells^15–17^. Yet, HLA class I is highly polymorphic (> 12,000 alleles), and the combinatorial peptide space exceeds 10^9^ sequences, making experimental identification of immunogenic candidates costly and laborious^18–20^.

Computational prediction of peptide–HLA binding has advanced substantially. Early methods such as SMMPMBEC^21^ (a Bayesian approach using pHLA binding energy covariance) and Consensus^22^ predictor built on the Immune Epitope Database (IEDB) improved affinity estimates. More recently, deep learning models, such as NetMHC 4.0^23,24^, MHCflurry 2.0^25^, TransPHLA^26^ and others^27–31^, have extracted complex sequence and structural features to enhance accuracy. Nevertheless, only a small fraction of top-ranked candidates exhibits true immunogenicity in validation assays^32^.

Recent generative algorithms aim to improve peptide selection by incorporating sampling strategies guided by those predictive models^26,33,34^. For example, AOMP integrates attention scores from TransPHLA^26^, while pepPPO^34^ employs reinforcement learning with MHCflurry 2.0-based^25^ rewards. However, reliance on existing prediction models can bias the search toward local regions of sequence space, limiting diversity and candidate quality.

To overcome these limitations, we developed AlphaVacc, a novel algorithm that directly generates optimized peptide datasets via reinforcement learning (RL)^35–38^, an approach that has powered AI breakthroughs such as AlphaGo Zero^38^ and DeepSeek-R1^39^. AlphaVacc rests on the hypothesis that randomly generated peptides generally lack immunogenicity, whereas peptides annotated as immunogenic in the IEDB retain their immunogenicity. We thus designed a peptide hallucination process combining Monte Carlo Tree Search^40,41^ (MCTS) and a Transformer^42^-based neural network. Through this hallucination process, AlphaVacc autonomously learns its own design logic to modify a peptide toward greater immunogenicity through supervised fine-tuning (SFT). In benchmarking tests across diverse peptide sources, AlphaVacc substantially outperformed previous generative models in identifying candidates with improved HLA binding affinity. Experimental validation of 12 candidates derived from a BING-4 protein template confirmed increased binding in 11 cases, and seven elicited significant T-cell responses. Further assessment of 16 single-point mutants validated 15 computational predictions. These results demonstrate that AlphaVacc efficiently produces high-potential candidates with enhanced immunogenicity and HLA affinity, offering a powerful computational platform for vaccine design.

## Results

### Architecture and training process of AlphaVacc

AlphaVacc is a deep reinforcement learning algorithm guided by MCTS for vaccine design (Fig. 1a). It modifies 9-mer antigen peptides—either patient-derived or randomly generated—to maximize immunogenicity for specific human leukocyte antigen (HLA) alleles. Starting from a peptide s, AlphaVacc iteratively proposes single-residue mutations using a 20 × 9 probability matrix derived from peptide hallucination process (details in Methods). Each hallucination cycle follows three MCTS stages: (1) *Selection*, in which the mutation *a*□ is chosen based on expected reward *Q* and exploration bonus *U*; (2) *Expansion* and *Evaluation*, in which a supervised fine-tuning network, SFT(AlphaVacc), is further introduced to predict both the mutation probability *p* and the success ratio *ν*; (3) and *Backup*, where *Q*-values and edge visit counts *N* are updated along the selected path (Fig. 1b). These three stages are then iterated over many cycles, allowing visit counts *N* to accumulate across all explored peptides. After a sufficient number of cycles, normalizing the visit counts *N* yields the probability distribution *π*(· |*s*), which guides the generation of novel neoantigen candidates.

**Figure 1.**
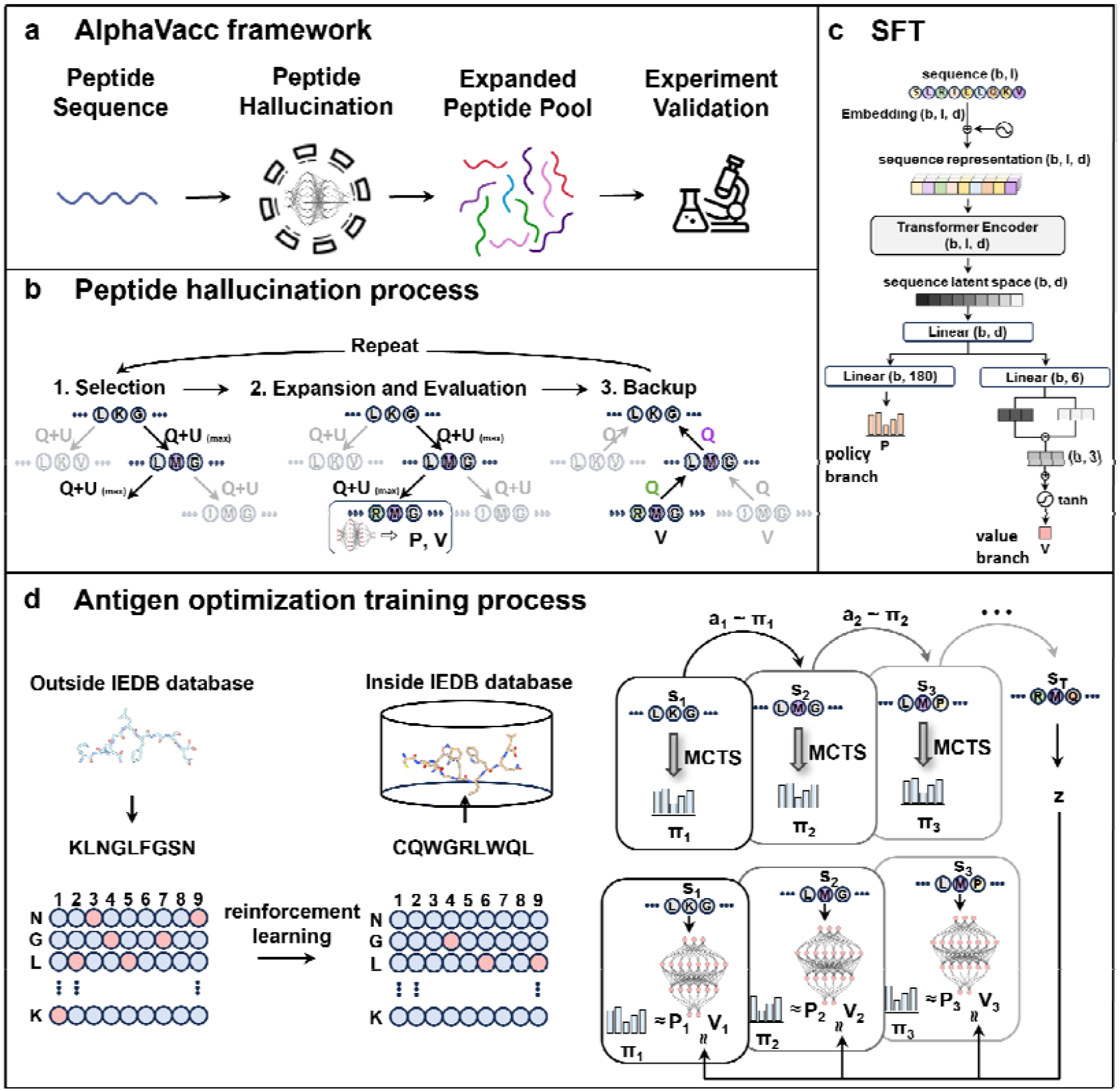
The AlphaVacc framework and training details. **(a)** AlphaVacc optimizes specific peptides through a peptide hallucination process that generates a high-quality expanded peptide pool serving as vaccine candidates for further experimental validations. **(b)** The peptide hallucination process, which combines MCTS and a Transformer-based neural network, comprises three main stages: Selection, Expansion and Evaluation, and Backup. **(c)** The structure of SFT(AlphaVacc) is illustrated. **(d)** Left: it is depicted how using reinforcement learning AlphaVacc optimizes a starting peptide sequence that is not in the IEDB database. The optimization process will be terminated when a sequence that is also in the IEDB database is generated or when reach the limited optimization steps. Right: the trajectory of AlphaVacc optimization process is shown. The sequences, and the corresponding probability matrices generated by hallucination process, and values obtained from the trajectory are collected to train a new SFT(AlphaVacc).

SFT(AlphaVacc), an embedded model within MCTS, is a Transformer-based supervised fine-tuning prediction network for peptide hallucination (Fig. 1c). It first converts input peptide sequences into embeddings, augments them with positional encodings, and then feeds them through a multi-head self-attention Transformer encoder to extract latent representations. Those representations then branch into two separate outputs: (1) the *policy* branch, consisting of linear layers producing a 1 × 180 vector *p* of residue mutation probabilities, and (2) the *value* branch, comprising linear layers with a *tanh* activation function that yields a scalar v representing the predicted success ratio.

The training of SFT(AlphaVacc) occurs in two stages: pretraining and fine-tuning. In the pretraining stage, we compile a binary classification dataset by retrieving pHLA binding pairs from IEDB for specific HLA phenotype. We train only the *value* branch using five-fold cross-validation to predict pHLA binding affinity. The *policy* branch is not pretrained and is initialized with Xavier uniform weights^43^. In the subsequent fine-tuning stage, both branches are jointly trained on a high-quality virtual dataset (Fig. 1d). We construct this dataset through iterative MCTS-based peptide hallucination: starting from a random peptide *s*_0_, we sample single-residue mutations *a*_*t*_ according to the probability distribution *π*(· |*s*_*t*_) produced by the hallucination process. We continue mutating until the current peptide matches any sequence in IEDB or until a preset step limit is reached. We assign *z* = 1 for successful matches (and *z* = 0 otherwise), and record each trajectory’s peptide sequences (*s*_0_,*s*_1_, …), the corresponding *π*(· |*s*_*t*_) at every step, and the final z value. These (*s, π, z*) tuples then serve as training examples to update both *policy* and *value* branches of SFT(AlphaVacc) (details in Arena competition process and Methods).

### Model evaluations

We evaluated the quality of peptides generated by AlphaVacc using four immunogenicity and HLA-binding predictors: Consensus^22^, SMMPMBEC^21^, MHCflurry 2.0^25^, and NetMHC 4.0^23^. These tools were used to assess (improved) “binder ratio”, the proportion of peptides predicted to show improved binding after optimization. For benchmarking, here we focused on peptides restricted to HLA-A*02:01 and considered two input types: biologically-derived and randomly-generated (Table 1). Each input group was optimized using AlphaVacc over 1,000 steps, with the process repeated across 10 independent runs. The resulting peptide sets were then evaluated and compared against those produced by baseline generative models, including Random, IEDB+BLOSUM62, TransPHLA^26^, and pepPPO^34^ (details in Methods).

**Table 1.**
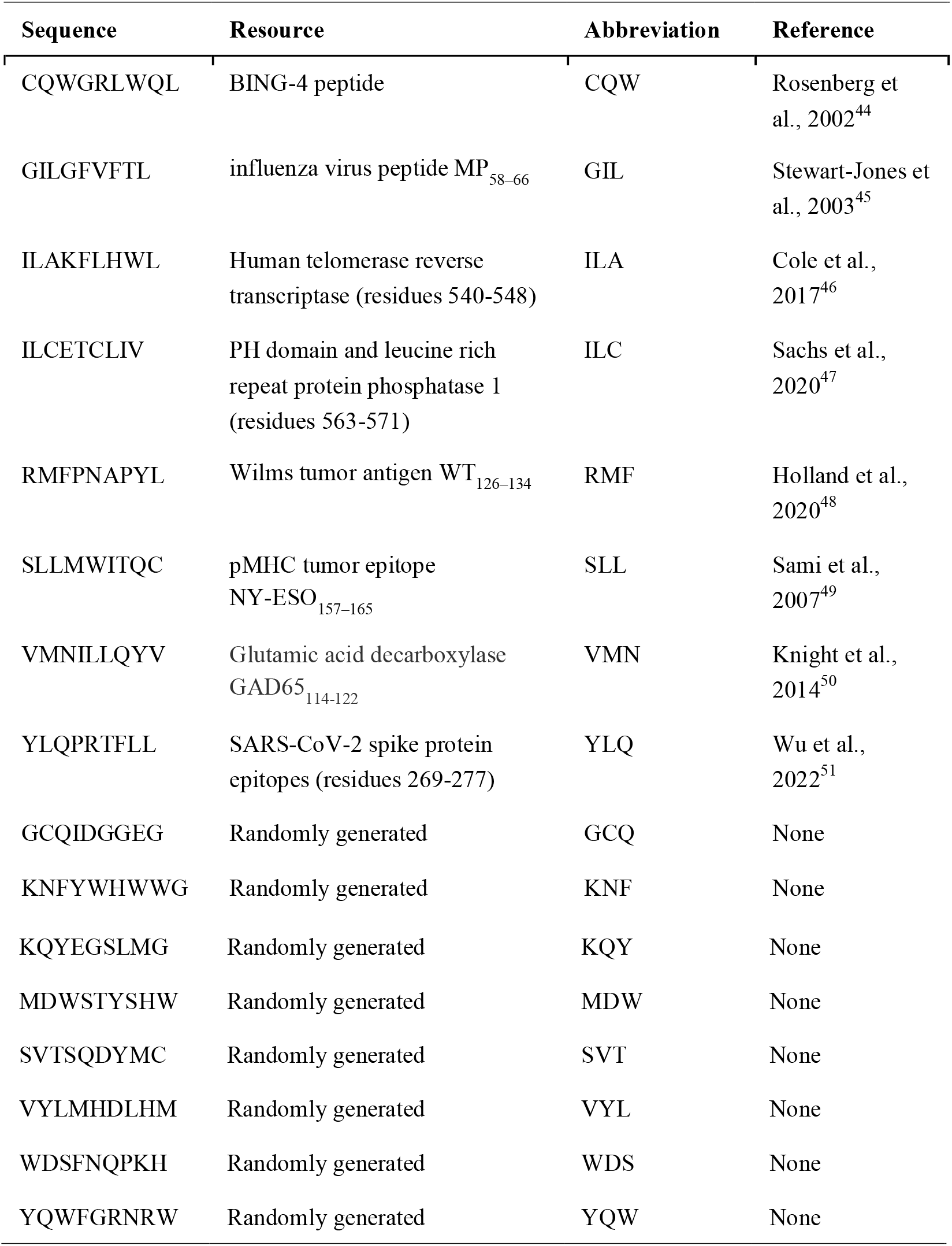
Biologically-derived peptides and randomly-generated peptides.

For biologically-derived peptides, AlphaVacc consistently outperformed all baseline generative models, achieving improved binder ratios exceeding 98% across most evaluation algorithms (Fig. 2a). SFT(AlphaVacc) alone also yielded acceptable performance, improving over 50% peptides, slightly below pepPPO. Notably, AlphaVacc produced a high sequence diversity (approximately 0.765), underscoring its capacity to explore a broad mutational space. When applied to randomly generated peptides, AlphaVacc achieved an even higher binder ratio of 99.6% (Fig. S1a).

**Figure 2.**
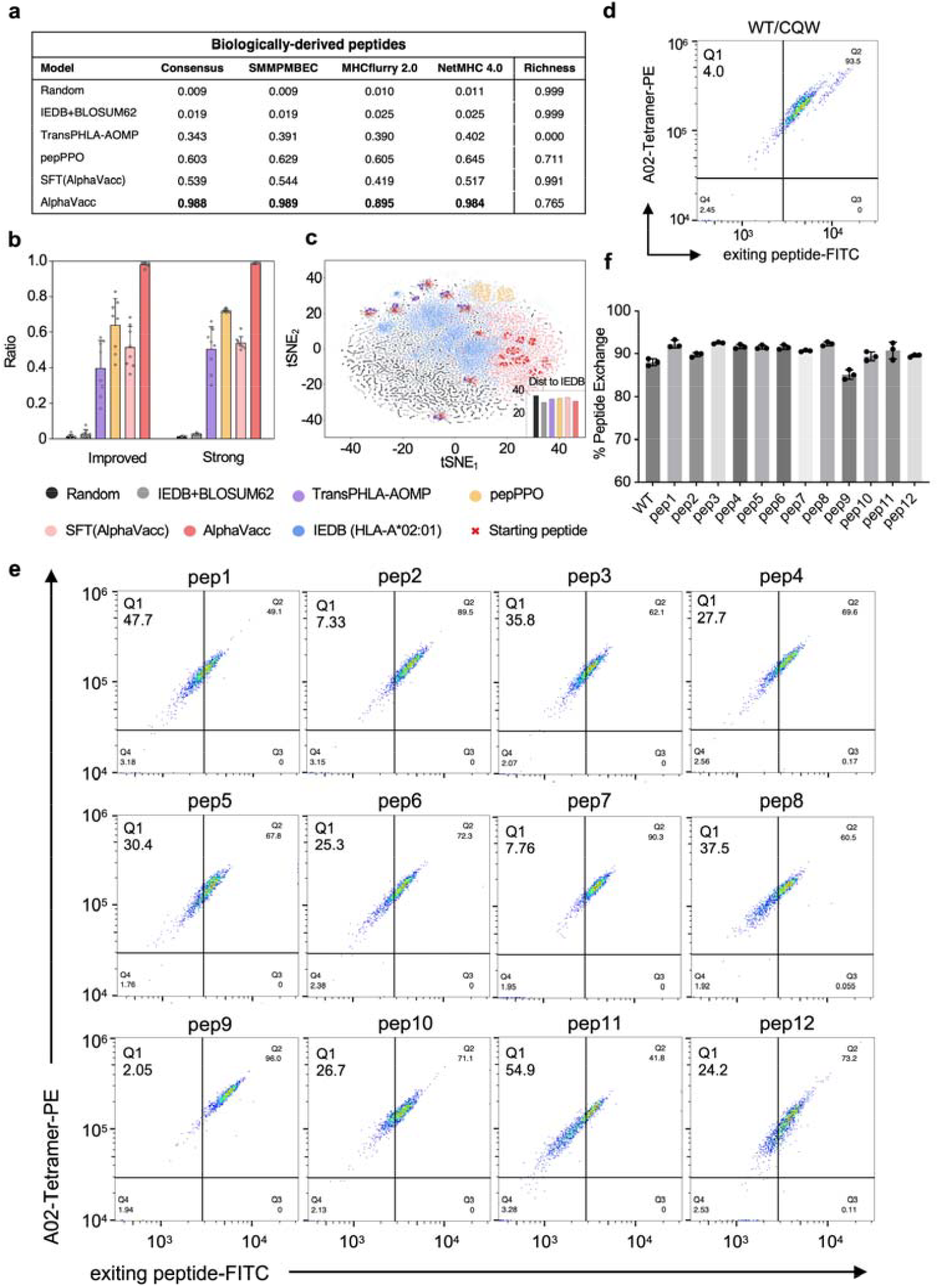
Peptide optimization quality comparison between AlphaVacc and other baseline models using biologically-derived peptides. **(a)** Different evaluation algorithms are used to assess the improved binder ratio in datasets generated by all the models. **(b)** A further comparison of the improved binder ratio and strong binder ratio of the different models. **(c)** The distribution of datasets generated by all the models, compared to that of the HLA-A*02:01 database in IEDB. The average Euclidean distances of different model datasets to IEDB are indicated in the bottom right corner. **(d)** Flow cytometry plots of the WT peptide. Gating strategy of flow cytometry was shown in Fig. S2. **(e)** Flow cytometry plots of the candidates. Q1 quadrant indicates the replacement percentage. **(f)** The peptide exchange efficiency of the WT peptide and all candidate peptides. The experiment was repeated three times. Data are shown as mean ± SD.

While generating improved binders is relatively straightforward for AlphaVacc, a more stringent benchmark is whether the optimized peptides qualify as “strong binders”, indicative of high immunogenic potential or binding affinity to HLA (details in Methods). For biologically derived peptides, AlphaVacc’s improved binder ratios were slightly lower than its strong binder ratios, though both exceeded those of baseline models (Fig. 2b). In contrast, for randomly generated peptides, AlphaVacc achieved markedly higher improved binder ratios while maintaining consistent strong binder performance (Fig. S1b). Notably, across all peptide sources, strong binder ratios exhibited smaller variance than improved binder ratios, suggesting greater robustness to input variability. AlphaVacc consistently maintained performance above 98% in both metrics, underscoring its effectiveness in generating high-quality vaccine candidates. Overall, it significantly outperformed all competing generative models.

To compare how each generative model samples peptide space relative to experimentally validated epitopes, we projected model-generated peptides and IEDB sequences into two dimensions using t-distributed stochastic neighbor embedding (t-SNE; details in Methods). The Random set dispersed most broadly, lying largely outside the principal IEDB cluster with highest mean Euclidean distance (Fig. 2c). In contrast, pepPPO remained tightly localized around the starting peptides, while IEDB+BLOSUM62, SFT(AlphaVacc) and AlphaVacc also extended into other IEDB regions. Consequently, each model occupied a distinct subspace of the peptide landscape. These spatial patterns were consistent regardless of whether the input peptides were biologically derived or randomly generated (Fig. S1c). Notably, the IEDB represents only a curated, high-confidence subset of immunogenic peptides, rather than the complete immunogenicity space. These results underscore the persistent challenge of systematically discovering immunogenic peptides.

To experimentally validate AlphaVacc, we optimized the BING-4 related peptide CQW (Table 1), referred here as the wild-type (WT) peptide to generate a pool of candidate peptides. From this pool, we selected 12 candidate peptides (Table 2) predicted to exhibit higher HLA-binding affinities than the WT. Their affinities were experimentally evaluated using a competitive peptide exchange assay, which quantifies how effectively test peptides displace a pre-bound peptide from HLA tetramers. Peptides with higher binding affinities are expected to more efficiently replace the pre-loaded peptide, resulting in greater exchange efficiency^52^. Assays were performed using the QuickSwitch Quant HLA-A*02:01 Tetramer Kit, which enables quantification of peptide exchange rates via flow cytometry (details in Methods). To quantify the peptide exchange efficiencies, we set quadrants on the flow cytometric plots based on the fluorescein FITC-conjugated mean fluorescence intensity (MFI_FITC_) of the WT peptide. The proportions of pHLA tetramers within each quadrant are labeled Q1, Q2, Q3, and Q4 (Fig. 2e). Flow cytometric analysis revealed that 11 of the 12 candidate peptides showed a leftward shift in MFI_FITC_ compared to the WT peptide. The change in Q1 values reflects the difference in MFI_FITC_ between peptides and serves as an indicator of replacement efficiency. According to the conversion formula provided by the kit, we used the MFI_FITC_ values to compute the peptide exchange efficiency. The results showed that while the exchange efficiency of WT was approximately 88%, 11 out of 12 candidates, exhibited superior exchange efficiencies compared to that of WT (Fig. 2f, Table S1). These results confirm that AlphaVacc is capable of efficiently generating optimized peptides with significantly enhanced MHC binding affinities.

**Table 2.**
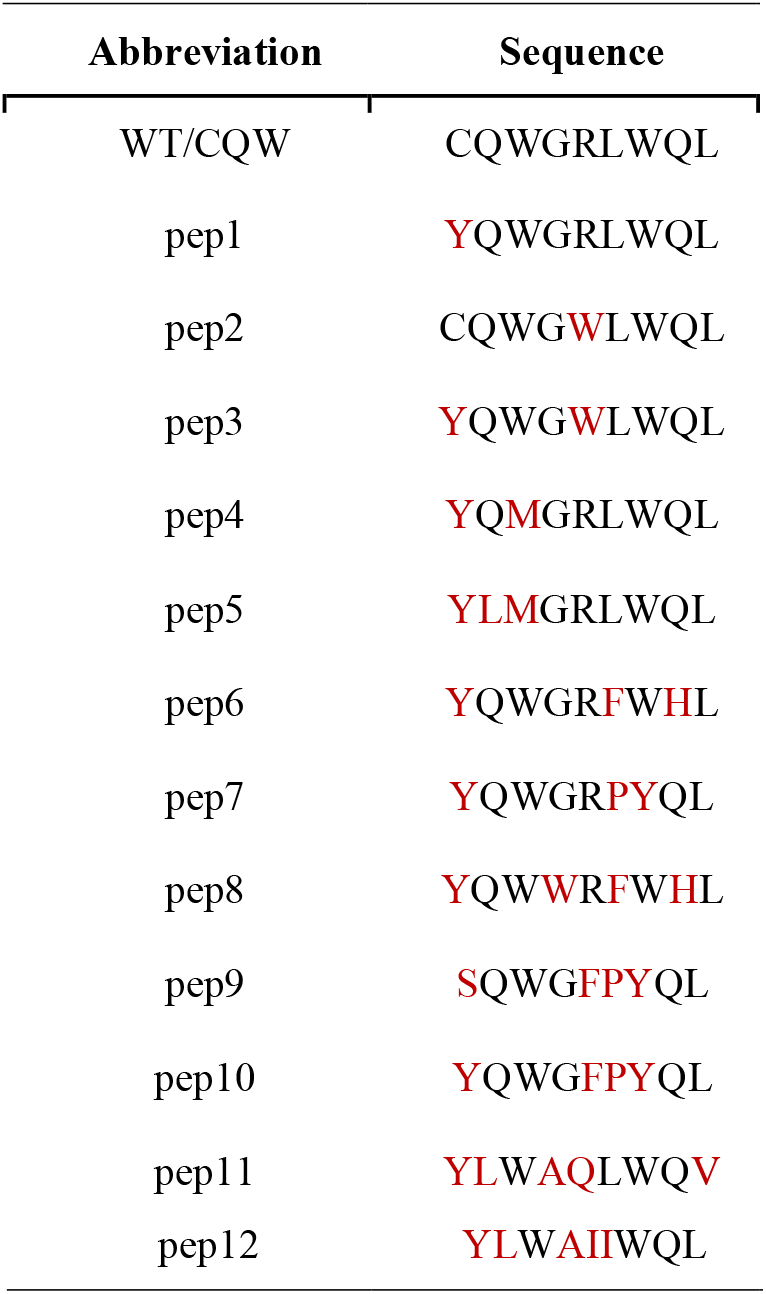
Amino acid sequences and abbreviations of candidate peptides from CQW.

### Immunogenicity assay of CQWGRLWQL neoantigen candidates

To further validate the immunogenicity potential of our candidate peptides, we assessed their ability to bind HLA molecules and induce immune responses using T2 binding assays and interferon-γ (IFN-γ) enzyme-linked immunosorbent spot (ELISpot) assays. The T2 binding assay evaluates how effectively peptides bind to HLA molecules presented by the T2 cell line (Fig. 3a), which is a human processing□defective cell line expressing empty HLA class I molecules. This system ensures exclusive presentation of exogenously loaded peptides^53^. Binding of peptides stabilizes HLA surface expression, which can be quantified through HLA staining followed by flow cytometry analysis (details in Methods). Our candidate peptides showed stable presentation on HLA*A-02:01 and exhibited stronger binding affinities than WT. These results were confirmed in duplicate T2 binding assays (Fig. 3c, Table S2).

**Figure 3.**
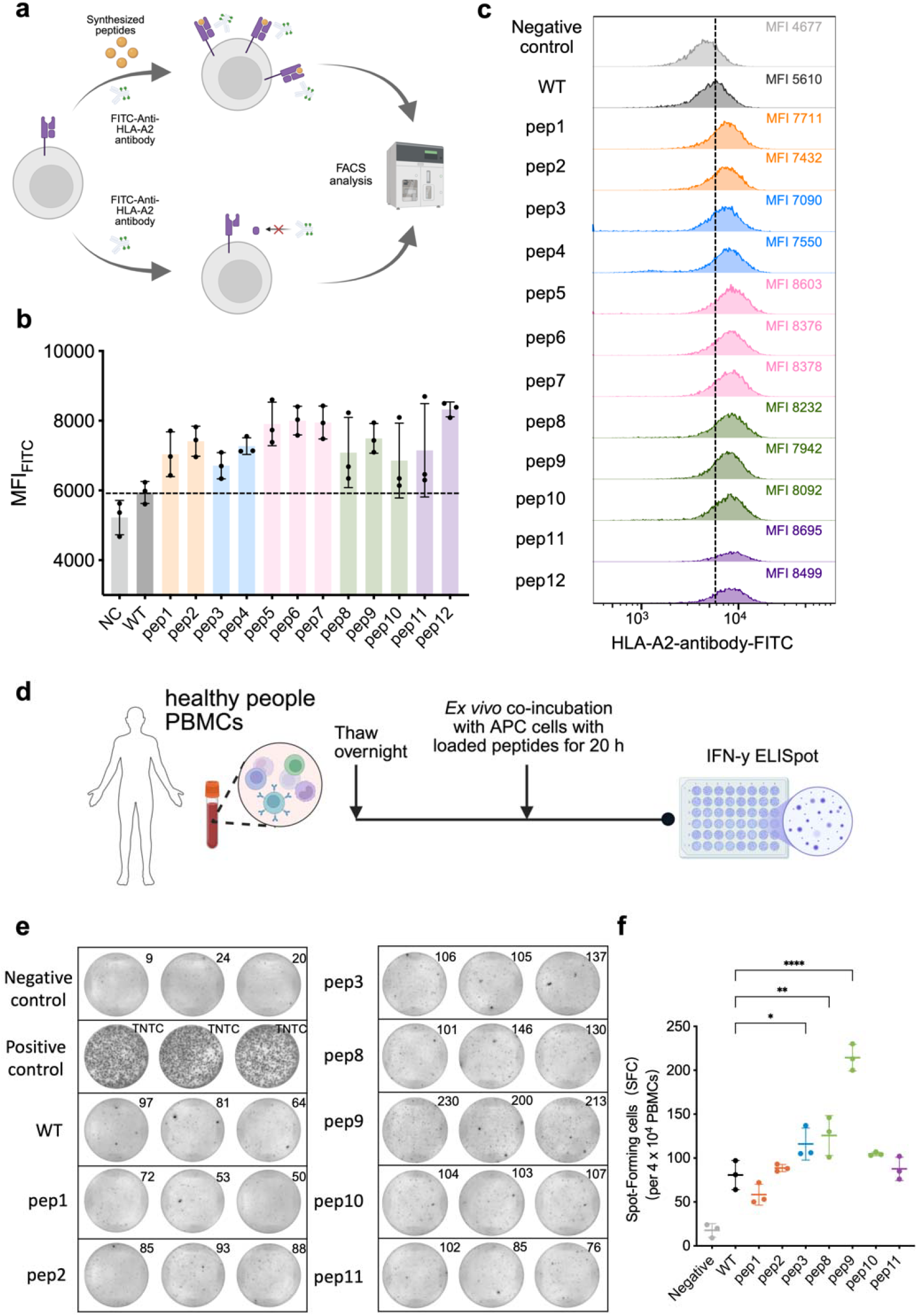
APC presentation and immunogenicity assay of neoantigen candidates. **(a)** Schematic diagram of the T2 peptide-binding assay. Created in https://BioRender.com. **(b)** The bar plot depicts the MFI_FITC_ of HLA-A*02:01 staining. The blue dashed line indicates the fluorescence intensity level of WT (n=3). **(c)** Representative flow cytometry plots of the T2 peptide-binding assays. Gating strategy of flow cytometry was shown in Fig. S3. The blue dashed line indicates the fluorescence intensity level of WT (n=3). **(d)** Schematic of the IFN-γ ELISpot assay. Created in https://BioRender.com. **(e)** Representative IFN-γ ELISpot images with HLA-A*02:01-positive homozygous donor cells. Only APCs and PMA/ionomycin treated served as negative and positive controls, respectively (n = 3). Only the peptides with responses similar or better than WT are shown. The numbers in the upper right corner of images represent spot counts. TNTC: Too Numerous To Count. **(f)** Statistical data for ELISpot data for this HLA-A*02:01-positive homozygous donor.

We next evaluated the ability of the candidate peptides to elicit immune responses using IFN-γ ELISpot assays, which detect secreted cytokines and count individual cytokine-producing cells^54^. IFN-γ, a key immunological marker, is predominantly secreted by activated CD4□ and CD8□ T cells, which together account for approximately 45–70% of peripheral blood mononuclear cells (PBMCs)^54^. PBMCs from different HLA-A*02:01-homozygous healthy donors were co-incubated with peptide-loaded antigen-presenting cells (APCs) (Fig. 3e, details in Methods). PBMCs cultured with APCs alone or in the presence of Phorbol 12-myristate 13-acetate (PMA) and ionomycin served as negative and positive controls, respectively. Among the 12 candidate peptides tested, three peptides (pep3, pep8, pep9) significantly enhanced human peripheral blood T-cell responses than WT, while four others induced comparable levels of T-cell activation (Fig. 3f, 3g, and S3a). In another donor assay, peptides pep1, pep2, pep7, pep8, pep9, and pep10 also triggered T-cell responses similar to the WT (Fig. S3b). Collectively, these results support the immunogenic potential of AlphaVacc-generated peptides and demonstrate its utility in identifying viable vaccine candidates.

### Arena competition process

The superior performance of AlphaVacc arises from the effective integration of MCTS with its supervised fine-tuning network, SFT(AlphaVacc). To obtain an effective SFT(AlphaVacc), we developed an iterative optimization framework termed “arena competition”, which systematically refines the model through data collection, fine-tuning, and direct performance comparison (Fig. 4a). This framework supports two training schemes, “full fine-tuning” and “partial fine-tuning”, denoted by the model prefix “F” and “P”, respectively (details in Methods). Each round of arena competition begins with the current best SFT(AlphaVacc) model embedded into the MCTS pipeline. This pipeline is used to optimize a batch of randomly initialized peptides, producing optimization trajectories that include peptide sequences *s*, mutation probability distributions *π*(*a*|*s*), and binary outcome indicators *z*. These newly generated trajectories form an expanded training set and are used to fine-tune a new SFT(AlphaVacc) model according to the selected training scheme. After fine-tuning, the newly trained model competes against the current best model by optimizing the same set of random peptides. The model that yields a greater number of successful outcomes (*z* = 1) is designated as the winner and promoted as the new best SFT(AlphaVacc) model for the next round. This cycle of data collection, model updating, and competitive evaluation enables continuous performance improvement through successive rounds of arena competition.

**Figure 4.**
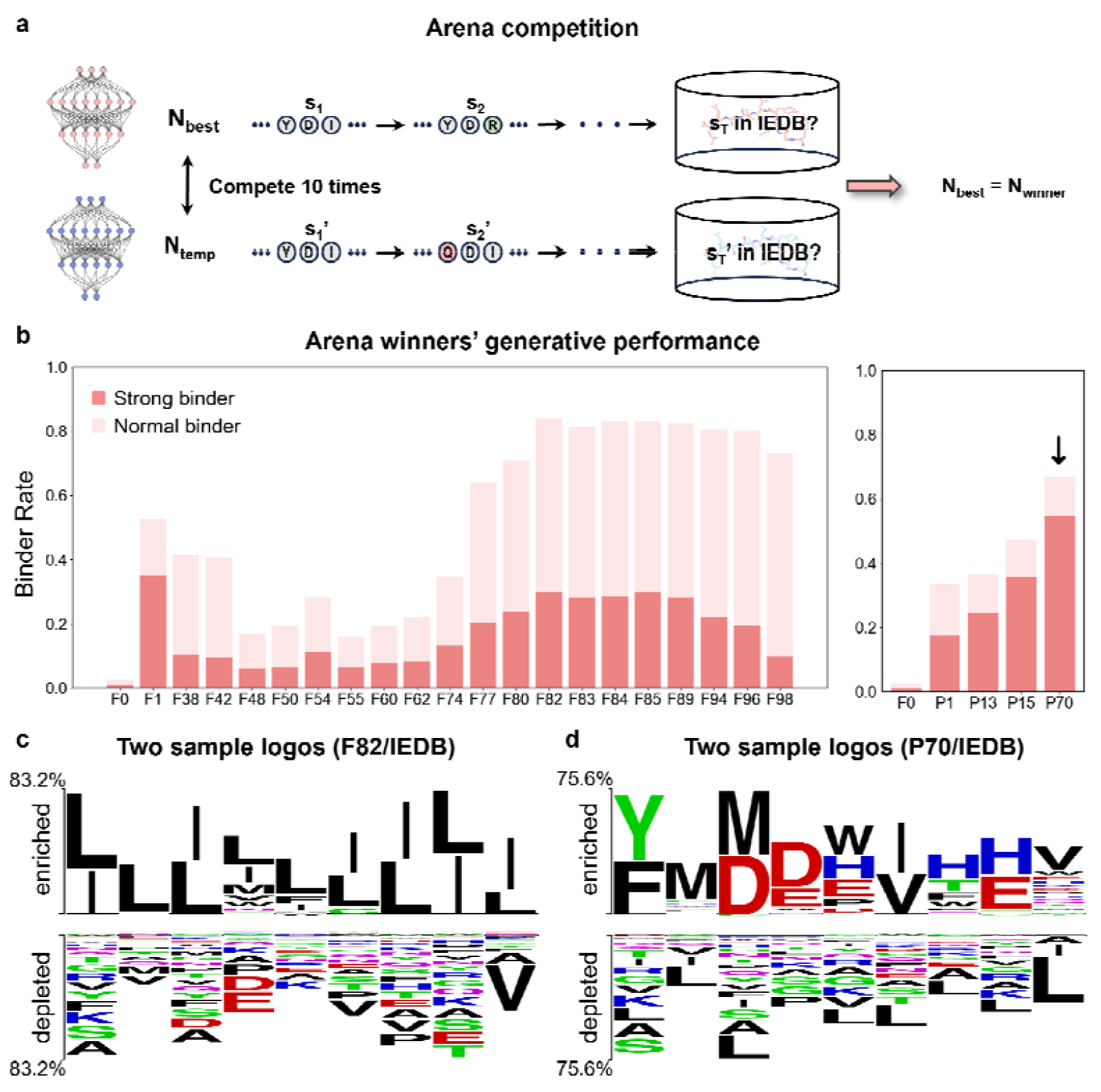
The arena competition process and analysis of its results. **(a)** A schematic representation of the arena competition between the current best model and a newly trained temporary model of SFT(AlphaVacc). They competed over 10 times, with the winner becoming the new best model. **(b)** Comparison of the ratios of the strong binders and normal binders among all the generated datasets of the arena winners, with both the full fine-tuning (F) and partial fine-tuning (P) networks. **(c)** Two sample logos generated from the datasets of ‘F82’ and IEDB. The ‘F82’ dataset is used as a positive sample while IEDB as a negative sample. **(d)** Two sample logos generated from the datasets of ‘P70’ and IEDB. The ‘P70’ dataset is used as a positive sample while IEDB as a negative sample. Symbol heights are proportional to the differences in symbol frequencies between the positive and negative samples. Enriched and depleted residues highlight the amino acids that are, respectively, over- or under-represented in the positive sample versus the negative sample. The percentage number in the y-axis shows the largest observed frequency difference between the two groups.

To visualize the performance improvements across arena competition rounds, we iteratively optimized a same peptide sequence using each model’s predicted probabilities. The resulting datasets were evaluated by calculating the proportions of normal and strong binders (Fig. 4b). Among all tested models, the ‘F82’ model achieved the highest combined binder ratio, while ‘P70’ produced the largest fraction of strong binders, over 50%. Overlap among optimized peptide sets remained below 21% across all models (Fig. S4a), indicating that each model explores distinct regions of the peptide sequence space.

We further compared amino acid preferences between peptides generated by different SFT(AlphaVacc) models and the IEDB target pool (Fig. 4d and S4b). The “F82” model, though delivered high combined binder ratio, exhibited signs of over-preference of hydrophobic residues, showing a strong bias toward those highly hydrophobic residues across all positions, which limited its sequence diversity. In contrast, ‘P70’ model displayed more balanced amino acid preferences, closely matching the experimental distributions. Specifically, leucine (L) and valine (V) were enriched at anchor positions 2 and 9, as well as at position 6, while aspartic acid (D) and glutamic acid (E) were favored at position 4, consistent with the reported experimental data^55^. Other positions also showed position-specific amino acid trends. These results suggest that partial fine-tuning mitigates potential “overfitting” (over-preference of certain type of amino acids) and enables the generation of peptides with both high quality and biologically realistic features. Overall, the arena competition process establishes a robust foundation for generating high-quality vaccine candidates.

### Ablation analysis

To isolate the contribution of MCTS to AlphaVacc’s performance, we conducted an ablation study involving four model variants: AlphaVacc_F.P._, SFT(AlphaVacc)_F.P._, AlphaVacc and SFT(AlphaVacc) (details in Methods). AlphaVacc_F.P._ and SFT(AlphaVacc)_F.P._ used fixed initial probabilities derived from AlphaVacc and SFT(AlphaVacc), respectively. All models were evaluated using four independent predictors, measuring improved binder ratios across the generated peptide sets. The results showed that removing MCTS reduced performance by over 30% (Fig. 5a), underscoring its important role in guiding efficient peptide optimization. Visualization of the generated peptide distributions using t-SNE (Fig. 5b) revealed that the peptide distribution from AlphaVacc_F.P._ were markedly distinct from those of the other models. Conversely, SFT(AlphaVacc)_F.P._, SFT(AlphaVacc), and AlphaVacc generated overlapping peptide distributions, with AlphaVacc producing a more focused cluster. This concentration suggests that the integration of MCTS enables AlphaVacc to effectively converge toward sequence features associated with strong immunogenicity, supporting its utility in vaccine candidate discovery.

**Figure 5.**
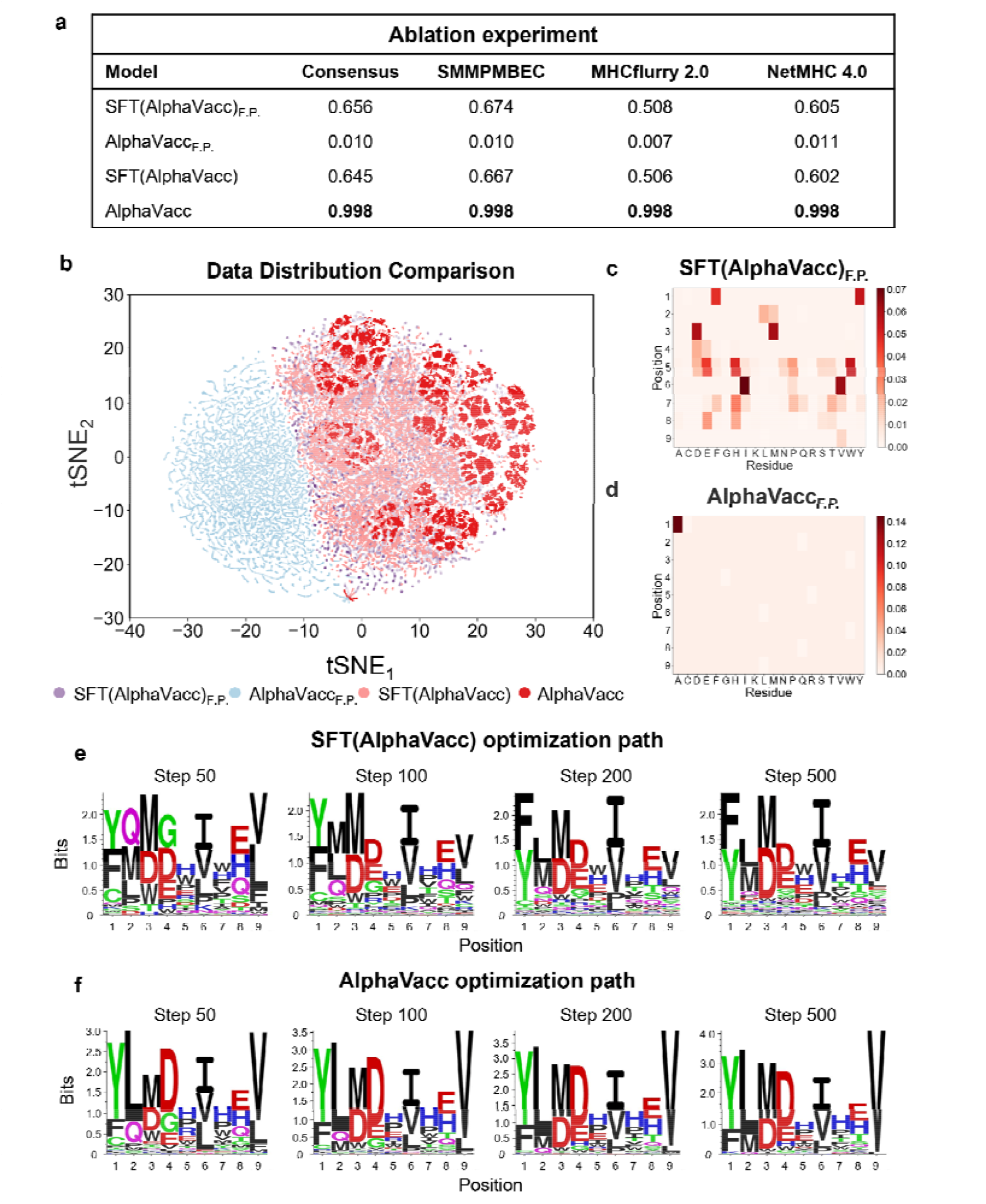
Ablation experiments validate the importance of MCTS performance in AlphaVacc. **(a)** To have deeper understanding of the performance of our model, we compared the four different peptide generation algorithms (AlphaVacc_F.P._, SFT(AlphaVacc)_F.P._, AlphaVacc and SFT(AlphaVacc)) derived from our model AlphaVacc. F.P. represents fixed initial probability. **(b)** Data distribution of the generated datasets by all these models. **(c)** The probability matrix of SFT(AlphaVacc)_F.P._. **(d)** The probability matrix of AlphaVacc_F.P._. **(e)** The computed motifs of the datasets generated by SFT(AlphaVacc) collected along the optimization path. **(f)** The computed motifs of the datasets generated by AlphaVacc collected along the optimization path.

Different model variants give rise to markedly distinct peptide generation behaviors, as reflected in their mutation preference profiles (Fig. 5c–f). The fixed probability matrix used by SFT(AlphaVacc)_F.P._ exhibited a relatively diverse and balanced amino acid distribution, while AlphaVacc_F.P._ showed a highly concentrated profile (Fig. 5c and 5d). This limited variability likely constrained its sequence exploration, contributing to its poor optimization performance. By comparison, AlphaVacc and SFT(AlphaVacc) alone dynamically updated their probabilities during the peptide optimization, enabling deeper exploration of the sequence space. The differences were further reflected in motif convergence behavior across ten independent optimization trials (Fig. 5e and 5f). AlphaVacc converged within ∼50 steps, closely matching the experimental motifs from IEDB (Fig. S5). In contrast, SFT(AlphaVacc) required around 200 steps to reach similar convergence. These results highlight that integrating MCTS not only enhances exploration efficiency but also accelerates convergence toward high-quality, immunogenic peptide candidates.

### Dual strategies captured by AlphaVacc and experimental validation

To trace the design logic behind AlphaVacc’s ability to generate strong binders, we focused on all optimized peptide sequences derived from the initial inputs in Table 1. This analysis aimed to reveal how AlphaVacc selects specific residues for mutation to enhance HLA binding affinity. We identified two dominant strategies employed by the model: a “*surge* strategy” and a “*progressive* strategy” (Fig. 6a). The choice of strategy depended on the initial affinity between the peptide and HLA. For peptides initially classified as poor binders, AlphaVacc adopted the *surge* strategy, prioritizing mutations at anchor positions 2 and 9 with hydrophobic residues (V, L, and methionine M). This behavior aligns with known preferences for HLA-A*02:01 anchor residues. In contrast, for peptides initially classified as normal binders, AlphaVacc applied the *progressive* strategy, introducing mutations at internal positions 3, 4, 5, 7, and 8 to incrementally improve binding affinity. Position 6 was rarely mutated, consistent with its hydrophobic and conserved nature. These findings illustrate how AlphaVacc dynamically adapts its mutation pathway to efficiently convert peptides into strong HLA binders.

**Figure 6.**
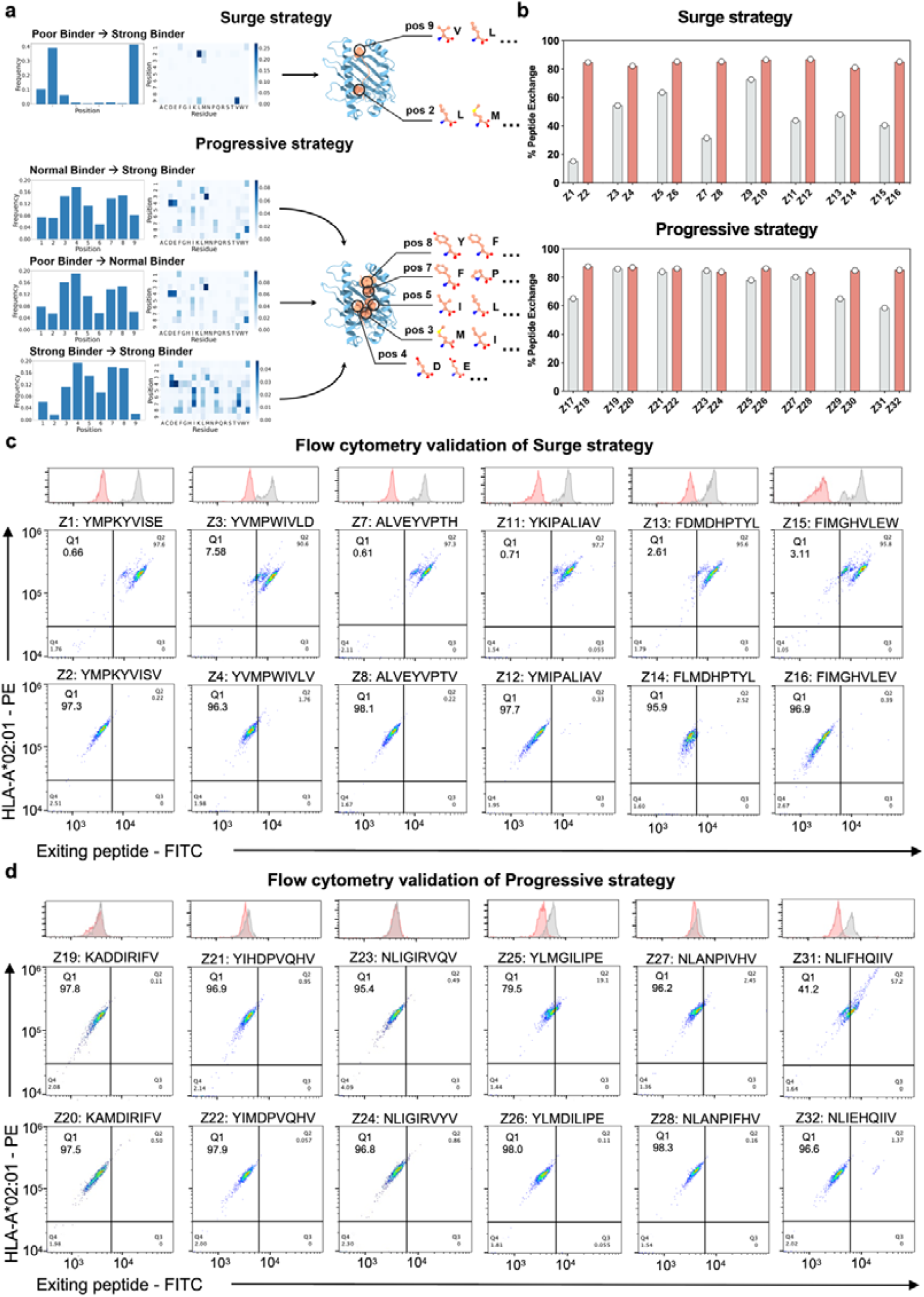
Two strategies from AlphaVacc design logic. **(a)** Two mutation strategies of AlphaVacc are illustrated: the *surge* strategy and the *progressive* strategy. **(b)** Peptide exchange efficiencies of the optimized peptides obtained by the *surge* and *progressive* strategies. **(c)** Representative flow cytometry plots of HLA-A*02:01 tetramer^+^ and the exiting peptide^-^ for validation of the *surge* strategy. **(d)** Representative flow cytometry plots of HLA-A*02:01 tetramer^+^ and exiting peptide^-^ for validation of the *progressive* strategy. Q1 quadrant indicates the peptide replacement percentage.

To experimentally validate these strategies, we selected 16 peptides, eight for each strategy, and assessed how single mutations influenced their HLA binding capacity (Table S3, S4). Peptide exchange assays (details in Methods) revealed that the peptides modified via the *surge* strategy showed significant improvements, transitioning from weak binding (under 60%) to strong binding (over 90%) after just one mutation. Similarly, peptides following *progressive* strategy showed enhanced binding when starting from normal or strong binders (Fig. 6b, Table S5). Flow cytometry (FCM) analysis further supported these findings, showing that optimized peptides displayed higher Q1 values compared to their original counterparts (Fig. 6c, 6d and S6). This indicates an increased competitive binding ability to HLA. Results from these two strategy groups indicate that out of the total 16 mutations, 15 agree with AlphaVacc’s predicted binding outcomes. This high success rate underscores the model’s capacity to capture meaningful, generalizable mutation patterns. Taking together, these results confirm that AlphaVacc can effectively evaluate the immunogenic potential of peptide, and select mutation strategies to drive efficient affinity optimization.

## Discussion

In this study, we introduced AlphaVacc, a generative framework that reengineers antigen optimization process through MCTS-guided peptide hallucination, which further includes SFT(AlphaVacc), a supervised fine-tunning neural network. Optimal neural network parameters are selected through an iterative arena competition process, ensuring robust performance. To evaluate its efficacy, we applied several established immunogenicity predictors. AlphaVacc consistently outperformed baseline generative models, achieving improved binder ratios exceeding 98% across multiple datasets. Most of these optimized peptides were also predicted to be strong binders, underscoring their immunogenic potential. As a first experimental validation, we optimized the BING-4 (CQW) peptide, generating 12 candidate vaccines. Eleven of these exhibited enhanced HLA binding affinity, and 7 induced significant T-cell responses. Ablation studies further demonstrated that integrating the MCTS algorithm was critical, improving peptide optimization performance by approximately 30% and enhancing the rationality of sequence modification decisions. Collectively, these results highlight AlphaVacc’s capacity to generate high-quality antigen sequences, supporting its potential utility in peptide vaccine design and broader immunotherapeutic applications.

To further elucidate AlphaVacc’s design logic, we analyzed the mutation trajectories leading to strong-binding peptides. Two dominant strategies emerged from this analysis: the *surge* strategy, applied primarily to peptides initially classified as poor binders, and the *progressive* strategy, used for peptides with moderate initial affinity. The *surge* strategy focused on anchor positions 2 and 9, favoring hydrophobic substitutions such as V and L, consistent with established HLA-A*02:01 binding motifs. In contrast, the *progressive* strategy selectively introduced mutations at internal positions, particularly 3, 4, 5, 7, and 8, to incrementally enhance binding affinity. Further experimental validations on 16 representative mutation groups showed strong agreement with AlphaVacc’s predictions, with 15 of the 16 groups confirming the expected binding outcomes. This high success rate underscores the model’s capacity to capture meaningful, generalizable mutation patterns. It is important to note, however, that this analysis focused specifically on transitions to stronger binders and does not fully capture AlphaVacc’s broader optimization dynamics across high-dimensional sequence space. Future studies could explore how AlphaVacc overcomes optimization barriers in peptides with limited improvement potential, for instance, how AlphaVacc overcomes optimization barriers for peptides with a limited improvement potential, or how it incorporates alternative evaluation criteria beyond binding affinity.

Despite its strong performance, AlphaVacc has certain limitations still. Although it effectively generates high-quality peptides, it does not fully recover all motif patterns present in the IEDB dataset. This gap highlights the potential benefit of incorporating external constraints or post-generation screening to guide peptide outputs toward desired amino acid distributions. Furthermore, for HLA alleles with limited or lower-quality training data, additional model refinement is likely necessary to achieve comparable results. To address these challenges, our further work will focus on optimizing the efficiency of peptide sampling method, adjusting the ratio of positive to negative samples in supervised fine-tuning datasets, and integrating tunable external constraints to enhance generalizability. We also aim to implement a more objective feedback mechanism within the reinforcement learning framework, combining data- and physics-driven criteria (e.g., surface physical/chemical metrics in the pHLA-TCR system). Ultimately, by incorporating additional biologically relevant information, we envision extending AlphaVacc into a unified, adaptable platform for vaccine design.

## Methods

### Peptide hallucination process

We repeatedly perform peptide hallucination for *N*_*self*_ iterations on the antigen peptides to generate a probability matrix. This 20×9-dimensional probability is a joint distribution probability, on which it is based to select the amino acids most likely to mutate. Each iteration of the hallucination process consists of three distinct stages: (1) Selection, (2) Expansion and Evaluation, and (3) Backup.

1. **Selection stage**: Starting from the root node (i.e. the current antigen peptide *s*_0_), the algorithm makes consecutive mutations. At each step t, the mutation action *a*_*t*_ is selected based on the equation:

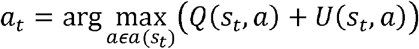

where *a*(*s*_*t*_) denotes the space of all possible mutation actions for the leaf node *s*_*t*_,*Q*(*s*_*t*_, *a*) is the average action value, and the exploration term *U*(*s*_*t*,_*a*) is calculated as:

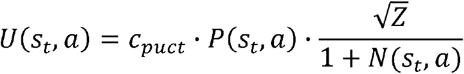 In this equation, *c*_*puct*_ is a parameter that characterizes the exploration breadth, *Z* represents the total visit number to the edge (*s*_*t*,_*b*) that takes every action *b*,

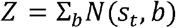 *P*(*s*_*t*_, *a*) denotes the prior probability of selecting the edge (*s*_*t*_, *a*), and (*s*_*t*_, *a*) is the number of times the edge (*s*_*t*_, *a*) is visited. This exploration process continues until it reaches the *L*-th step or encounters a previously unvisited leaf node, either case denoted as *S*_*L*_, at which point the exploration stops.
2. **Expansion and Evaluation stage**: Upon reaching a new leaf node *S*_*L*_, we apply the SFT(AlphaVacc) neural network, denoted as *f*_*θ*_ (·), to predict the amino acid mutation probability, *p*_*net*_(*S*_*L*_)and the target success rate *ν*_*net*_ (*S*_*L*_):

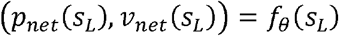 At the same time, we initialize the statistical values for each possible action (*s*_*L*_, *a*) from node *S*_*L*_:

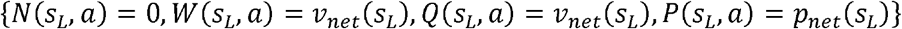 Here, *N*(*s*_*L*_, *a*) represents the number of times the edge (*s*_*L*_, *a*) is visited; *W*(*s*_*L*_, *a*) represents the total action value, *Q* (*s*_*L*_, *a*) represents the average action value; and *P*(*s*_*L*_, *a*) represents the prior probability of selecting the edge (*s*_*L*_, *a*).
3. **Backup stage**: During this stage, we update the statistical information for each edge encountered in the selection stage. Specifically, we increment the visitation count:

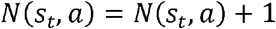

and the total action value and the average action value as follow:

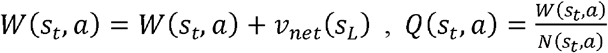 After performing the above three stages for *N*_*self*_ iterations, we compute the final probability matrix as:

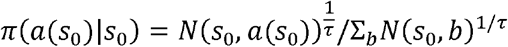

where *τ* is a temperature parameter controlling the extent of exploration; and *a*(*s*_0_) represents the space of all possible mutation actions for the root node *s*_0_. This probability matrix *π* will be used as probability distribution in the numpy.random.choice function to select the next single mutation of the peptide sequence. Additionally, we constrain this new peptide sequence to be the one that we never met before.

### SFT(AlphaVacc) network

SFT(AlphaVacc) is a Transformer-based prediction network that treats each residue of the epitope sequence as an individual input token. A special [CLS] token is prepended to each sequence to capture its overall features, resulting in a vocabulary consisting of 20 amino acids plus the [CLS] token. These tokens are processed by an embedding layer and subsequently enriched with positional information via a positional encoding layer. Following this, a Transformer encoder consisting of multi-head self-attention layers learns latent representations. The latent representation corresponding to the [CLS] token is then branched into two separated paths: the *policy* branch, where linear layers produce a vector with predicted amino acid mutation probability, *p*_*net*_(*S*_*L*_), and the *value* branch, where linear layers combined with a *tanh* activation function generate a scalar representing the predicted success rate *ν* for achieving the target.

### Pretraining process

We collected and curated a dataset containing 57,340 unique 9-mer peptides from the IEDB database, labeled according with their ability to be presented by HLA-A*02:01. This refined dataset is denoted as the IEDB binary dataset, utilized for five-folds cross-validation training during the pretraining phase. This pretraining process only targets the *value* branch, optimizing the weights of SFT(AlphaVacc) without the *policy* branch. The weights of the *policy* branch in SFT(AlphaVacc) will be initialized using a Xavier uniform distribution for further supervised fine-tuning processes.

### Supervised fine-tuning process

We implemented two fine-tuning strategies for our network: full fine-tuning and partial fine-tuning. In the former approach, all neural network weights are optimized through backpropagation. In contrast, the latter approach optimizes only the linear layers following the Transformer encoder, while all other weights remain unchanged after pretraining.

During the fine-tuning, both *policy* and *value* branches of SFT(AlphaVacc) are simultaneously trained using a high-quality virtual database. We assume that peptides from the IEDB database represent a high level of immunogenicity due to their extensive experimental validation, and thus we use these peptides as our target pool. Our goal is to iteratively modify a randomly initialized peptide using the probability matrix *π* generated from the peptide hallucination process, until the modified peptide matches a sequence in the target pool. Specifically, each optimization round starts with *N*_*iter*_=100 randomly initialized antigen peptides. Within each round, we apply two levels of success control: a “soft” and “rigorous” success check. The soft success check, used for the first 90% of the optimization attempts, is met if SFT(AlphaVacc) predicts a success rate above 0.95 within *N*_*episoed*_=1000 steps. The rigorous success check, employed for the remaining 10% of attempts, requires an exact sequence match with an antigen peptide from the IEDB database. The combination of the criteria balances its computational efficiency with its optimization accuracy. Along each optimization trajectory, we record the optimized sequences s, the corresponding probability matrices *π*(*a* | *s*)derived from the peptide hallucination process, and the binary vales *z*, where z=1 if the target criterion is achieved within *N*_*episoed*_ steps and z=-1 otherwise.

The generated virtual database (*s*,(*π* (*a* | *s*),*z*)) used for the supervised fine-tuning consists of the most recent 500 optimization trajectories, as larger databases would significantly increase training time. The loss function used in this process is defined as:

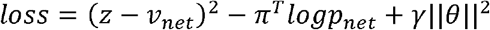

where *θ* denotes the neural network parameters. This approach allows us to obtain multiple trained models from which we select the best-performing version.

### Evaluation algorithms

Evaluation algorithms are used to assess the generated peptide quality. NetMHC 4.0 serves as our primary evaluation standard, harnessing its sequence alignment capabilities based on artificial neural networks (ANN). Peptides are classified based on their rank values: those with a rank less than 0.5 are considered strong binders, those ranked between 0.5 and 2 as normal binders, and those ranked greater than 2 as poor binders. Furthermore, we utilize other predictive scoring functions, such as MHCflurry 2.0, SMMPMBEC, and Consensus. All these methods are used to evaluate the proportion of improved binders among all the generated peptides to get the improved binder ratio.

### Baseline models

To evaluate the quality of the generated peptides by AlphaVacc and SFT(AlphaVacc), we compare them against the peptides produced by four baseline models: Random, IEDB+BLOSUM, TransPHLA-AOMP, and pepPPO. The Random and IEDB+BLOSUM models do not incorporate structural or MHC interaction information. Specifically, the Random model generates a new peptide pool by starting from a reference peptide and applying a uniform distribution probability to select both the mutation positions and amino acid residues. In contrast, the IEDB+BLOSUM model uses a uniform distribution probability to select a mutation position but applies a combination of equally weighted BLOSUM62 frequency and IEDB frequency (derived from the Immune Epitope Database, IEDB), to determine the residue mutations. The TransPHLA-AOMP model utilizes the TransMut framework to optimize peptide mutations guided by attention scores from the transformer-based model, TransPHLA. The pepPPO model represents another innovation, using reinforcement learning to generate target peptides for specific MHC alleles, reinforcing its selections using predictions from the MHCflurry2.0 algorithm.

### Sequence logo

We used sequence logos to visually represents the information content and significance of peptide sequence positions. A sequence logo graphically displays amino acid frequencies at each position in the peptide sequence. Each position is represented by a stack of letters, where the height of each letter reflects its relative frequency in the sequences. The total height at each position corresponds to the position’s overall information content, measured in bits. Specifically, the information content *I*_*j*_ of position *j* is given by:

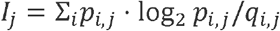

Where *p*_*i,j*_ represents the observed probability (calculated from the database) of amino acid *i* in position *j*, and *q*_*i,j*_ represents its background probability. Using Shannon’s sequence logo approach, the height of amino acid letter *i* in *j* position is:

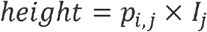

where a taller letter indicates a higher relative frequency of that amino acid at that position.

### Two Sample Logo

Sequence logos cannot easily visualize the differences between two groups of aligned sequences. We use Two Sample Logos (TSL)^56^ to calculate and visualize the differences between the generated datasets of the different versions of SFT(AlphaVacc) and IEDB. This approach visualizes the statistically significant differences in amino acid occurrences between two datasets of aligned sequences. In our work, peptides generated by different SFT(AlphaVacc) versions (‘F82’ and ‘P70’) constitute the positive sample, while the IEDB peptides serve as the negative sample, facilitating a clear comparison of the dataset differences. The upper or lower section displays the symbols enriched (overrepresented) or depleted (underrepresented) in the positive set.

### Distribution Comparison through the t-SNE analysis

We apply t-distributed Stochastic Neighbor Embedding (t-SNE), a statistical method for visualizing high-dimensional data in lower dimensions, to compare data distributions between the target IEDB pool and peptide sequences generated during model evaluation. This method allows us to map each datapoint into a two- or three-dimensional map. All the generated data were delivered from six models (Random, IEDB_BLOSUM62, pepPPO, TransPHLA-AOMP, SFT(AlphaVacc), and AlphaVacc), under both biologically-derived and randomly-generated conditions. Additionally, four models (AlphaVacc_F.P._, SFT(AlphaVacc)_F.P._, AlphaVacc and SFT(AlphaVacc)) used in the ablation study were analyzed to interpret their relationship.

All peptide sequences were transformed into high-dimensional vectors through one-hot encoding, representing each amino acid position with 21 features, including a placeholder “X” for any non-standard residue. The encoding is expressed as:

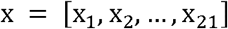

where each *x*_*i*_ represents a binary feature indicating the presence or absence of the *ith* amino acid in the sequence position.

The high-dimensional data was then visualized in two dimensions using t-SNE, which minimizes the divergence between high-dimensional and low-dimensional probability distributions. This divergence is given by the Kullback-Leibler (KL) divergence:

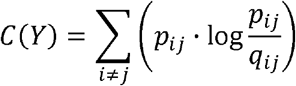

where *p*_*ij*_ is the pairwise probability distribution in the high-dimensional space, and *q*_*ij*_ is the corresponding pairwise probability distribution in the low-dimensional space.

### Peptide-HLA tetramer exchange assays

To evaluate the binding affinities of peptides to HLA, we used the QuickSwitch Quant HLA-A*02:01 Tetramer kit (PE-labeled, Cat. No. TB-7300-K1, MBL International), and performed the peptide exchange assays following the manufacturer’s protocol.

Briefly, all peptides were chemically synthesized (Sangon Biotech, Shanghai, China) and dissolved in DMSO to prepare a 1 mM stock solution. Each of the peptide was mixed with QuickSwitch™ Tetramer in a well containing the Magnetic Capture Beads, followed by the Exiting Peptide Antibody-FITC. Three controls were set up: Control #1 was a negative control that omitted the Exiting Peptide Antibody-FITC or any test peptide; Control #2 was another negative control that omitted the tetramers whereas Control #3 was a positive control that was mixed with the Exiting Peptide Antibody-FITC only without any test peptide. Flow cytometry was used to measure the mean fluorescence intensity (MFI) of the fluorescein FITC from the singlet beads after its gating parameters were optimized using the FlowJo™ Software (Fig. S2 and S6) such that the bead doublets and aggregates were excluded. The measured MFI_FITC_ values of test peptides fall between that of Controls #2 (0% Exiting Peptide Antibody-FITC) and #3 (100% Exiting Peptide Antibody-FITC). And the level of Exiting Peptide-Peptide Antibody-FITC is inversely proportional to the amount of the newly loaded peptides on the MHC molecules.

### T2 binding assay

The binding capacity of peptides to HLA-A*0201 molecules was measured using T2 cells, that is, a TAP-deficient, HLA-A*0201-positive human cell line. Briefly, T2 cells were washed three times, resuspended at a density of 10^6^ cells/ml, and incubated at 37 ºC for 1.5 hours in a serum-free medium containing 80 µM of each peptide. After incubation, cells were washed twice with the cold FACS buffer (PBS with 2% FCS). FITC-conjugated mouse anti-HLA-A2 antibody (Cat# 343304, BioLegend) was added to the cell suspension at 100 µL/10^6^ cells and incubated at 4ºC in the dark for 20 min. Cells were then washed twice with the cold FACS buffer. HLA-A2 surface expression was analyzed using fluorescence-activated cell sorter scan (FACS, Sony SH800S, Japan). All T2 binding assays were carried out in duplicates.

### ELISpot assay

Peripheral blood mononuclear cells (PBMCs) from healthy donors were purchased from Milecell Biological Science & Technology Co., Ltd. IFN-γ ELISpot assays were performed using the PBMCs from healthy human donors according to the manufacturer’s protocol (Cat. ab62899, Abcam). PBMCs (4×10^4^ cells per well) were co-incubated with the peptide-loaded antigen-presenting cells (APCs, 2×10^5^ cells per well) for 20 hours using a human IFN-γ pre-coated ELISpot kit, following the standard protocol (Cat. ab62899, Abcam). In parallel, PBMCs were cultured with APCs alone as the negative control, or with Phorbol 12-myristate 13-acetate (PMA, Cat. 16561-29-8, MCE) and ionomycin (Cat. 56092-81-0, MCE) as positive controls. After incubation, ELISpot plates were air-dried and imaged using a stereo fluorescence microscope (Nikon SMZ18). Images presented in Fig. 3 and S3 were converted to grayscale for clarity. Brightness and contrast adjustments were applied uniformly across all images for each donor. ELISpot plates were counted using the ImageJ software.

## Supporting information

Cover letter

Supplementary Information-AlphaVacc-bioRxiv

## Data availability

The data used in this article are publicly available at https://github.com/ZGQVictory/AlphaVacc/tree/master. All code used in this study and the final trained models are provided in our public GitHub repository: https://github.com/ZGQVictory/AlphaVacc/tree/master. Users could update AlphaVacc to other situations related to peptide generation by their data using our training procedure.

## Acknowledgments

This work was partially supported by the National Key R&D Program of China (2024YFA1306400, 2021YFA1201200, 2024YFA1307500), the National Natural Science Foundation of China (U1967217), the National Center of Technology Innovation for Biopharmaceuticals (NCTIB2022HS02010), Shanghai Artificial Intelligence Lab (P22KN00272), the National Independent Innovation Demonstration Zone Shanghai Zhangjiang Major Projects (ZJZX2020014), the Starry Night Science Fund of Zhejiang University Shanghai Institute for Advanced Study (SN-ZJU-SIAS-003), and Zhejiang University Global Partnership Fund (188170+194452505).

## References

1. Bray, F. et al. Global cancer statistics 2022: GLOBOCAN estimates of incidence and mortality worldwide for 36 cancers in 185 countries. CA: A Cancer Journal for Clinicians 74, 229–263 (2024).

2. Chen, S. et al. Estimates and Projections of the Global Economic Cost of 29 Cancers in 204 Countries and Territories From 2020 to 2050. JAMA Oncology 9, 465–472 (2023).

3. Yaremenko, A. V., Khan, M. M., Zhen, X., Tang, Y. & Tao, W. Clinical advances of mRNA vaccines for cancer immunotherapy. Med 6, (2025).

4. Ikeda, H. Cancer immunotherapy in progress—an overview of the past 130 years. International Immunology dxaf002 (2025) doi:10.1093/intimm/dxaf002.

5. Varadé, J., Magadán, S. & González-Fernández, Á. Human immunology and immunotherapy: main achievements and challenges. Cell Mol Immunol 18, 805–828 (2021).

6. Waldman, A. D., Fritz, J. M. & Lenardo, M. J. A guide to cancer immunotherapy: from T cell basic science to clinical practice. Nat Rev Immunol 20, 651–668 (2020).

7. Haslam, A. & Prasad, V. Estimation of the Percentage of US Patients With Cancer Who Are Eligible for and Respond to Checkpoint Inhibitor Immunotherapy Drugs. JAMA Network Open 2, e192535 (2019).

8. Research, C. for D. E. and. FDA grants accelerated approval to afamitresgene autoleucel for unresectable or metastatic synovial sarcoma. FDA (2024).

9. Lin, M. J. et al. Cancer vaccines: the next immunotherapy frontier. Nat Cancer 3, 911–926 (2022).

10. Hu, Z., Ott, P. A. & Wu, C. J. Towards personalized, tumour-specific, therapeutic vaccines for cancer. Nat Rev Immunol 18, 168–182 (2018).

11. Fan, T. et al. Therapeutic cancer vaccines: advancements, challenges and prospects. Sig Transduct Target Ther 8, 1–23 (2023).

12. Liu, J. et al. Cancer vaccines as promising immuno-therapeutics: platforms and current progress. J Hematol Oncol 15, 28 (2022).

13. Zhang, Z. et al. Neoantigen: A New Breakthrough in Tumor Immunotherapy. Front. Immunol. 12, (2021).

14. Jou, J., Harrington, K. J., Zocca, M.-B., Ehrnrooth, E. & Cohen, E. E. W. The Changing Landscape of Therapeutic Cancer Vaccines—Novel Platforms and Neoantigen Identification. Clinical Cancer Research 27, 689–703 (2021).

15. Minati, R., Perreault, C. & Thibault, P. A Roadmap Toward the Definition of Actionable Tumor-Specific Antigens. Frontiers in Immunology 11, (2020).

16. Jensen, P. E. Recent advances in antigen processing and presentation. Nat Immunol 8, 1041–1048 (2007).

17. Pishesha, N., Harmand, T. J. & Ploegh, H. L. A guide to antigen processing and presentation. Nat Rev Immunol 22, 751–764 (2022).

18. Sarkizova, S. et al. A large peptidome dataset improves HLA class I epitope prediction across most of the human population. Nat Biotechnol 38, 199–209 (2020).

19. Luescher, I. F., Romero, P., Cerottini, J. C. & Maryanski, J. L. Specific binding of antigenic peptides to cell-associated MHC class I molecules. Nature 351, 72–74 (1991).

20. Elvin, J., Cerundolo, V., Elliott, T. & Townsend, A. A quantitative assay of peptide-dependent class I assembly. Eur J Immunol 21, 2025–2031 (1991).

21. Kim, Y., Sidney, J., Pinilla, C., Sette, A. & Peters, B. Derivation of an amino acid similarity matrix for peptide:MHC binding and its application as a Bayesian prior. BMC Bioinformatics 10, 394 (2009).

22. Moutaftsi, M. et al. A consensus epitope prediction approach identifies the breadth of murine TCD8+-cell responses to vaccinia virus. Nat Biotechnol 24, 817–819 (2006).

23. Andreatta, M. & Nielsen, M. Gapped sequence alignment using artificial neural networks: application to the MHC class I system. Bioinformatics 32, 511–517 (2016).

24. Nielsen, M. et al. Reliable prediction of T-cell epitopes using neural networks with novel sequence representations. Protein Science 12, 1007–1017 (2003).

25. O’Donnell, T. J., Rubinsteyn, A. & Laserson, U. MHCflurry 2.0: Improved Pan-Allele Prediction of MHC Class I-Presented Peptides by Incorporating Antigen Processing. Cell Systems 11, 42-48.e7 (2020).

26. Chu, Y. et al. A transformer-based model to predict peptide–HLA class I binding and optimize mutated peptides for vaccine design. Nat Mach Intell 4, 300–311 (2022).

27. Lu, T. et al. Deep learning-based prediction of the T cell receptor–antigen binding specificity. Nat Mach Intell 3, 864–875 (2021).

28. Springer, I., Tickotsky, N. & Louzoun, Y. Contribution of T Cell Receptor Alpha and Beta CDR3, MHC Typing, V and J Genes to Peptide Binding Prediction. Front. Immunol. 12, (2021).

29. Yu, C., Fang, X., Tian, S. & Liu, H. A unified cross-attention model for predicting antigen binding specificity to both HLA and TCR molecules. Nat Mach Intell 7, 278–292 (2025).

30. Feng, Z. et al. Sliding-attention transformer neural architecture for predicting T cell receptor–antigen–human leucocyte antigen binding. Nat Mach Intell 6, 1216–1230 (2024).

31. Wohlwend, J. et al. Deep learning enhances the prediction of HLA class I-presented CD8+ T cell epitopes in foreign pathogens. Nat Mach Intell 7, 232–243 (2025).

32. Nibeyro, G. et al. Unraveling tumor specific neoantigen immunogenicity prediction: a comprehensive analysis. Front. Immunol. 14, (2023).

33. Lin, Y. et al. Integrating Reinforcement Learning and Monte Carlo Tree Search for enhanced neoantigen vaccine design. Briefings in Bioinformatics 25, bbae247 (2024).

34. Chen, Z. et al. Binding peptide generation for MHC Class I proteins with deep reinforcement learning. Bioinformatics 39, btad055 (2023).

35. Kiran, B. R. et al. Deep Reinforcement Learning for Autonomous Driving: A Survey. IEEE Transactions on Intelligent Transportation Systems 23, 4909–4926 (2022).

36. Silver, D. et al. Mastering the game of Go with deep neural networks and tree search. Nature 529, 484–489 (2016).

37. Fawzi, A. et al. Discovering faster matrix multiplication algorithms with reinforcement learning. Nature 610, 47–53 (2022).

38. Silver, D. et al. Mastering the game of Go without human knowledge. Nature 550, 354–359 (2017).

39. DeepSeek-AI et al. DeepSeek-R1: Incentivizing Reasoning Capability in LLMs via Reinforcement Learning. Preprint at 10.48550/arXiv.2501.12948 (2025).

40. Coulom, R. Efficient Selectivity and Backup Operators in Monte-Carlo Tree Search. In Computers and Games (eds. van den Herik, H. J., Ciancarini, P. & Donkers, H.H.L.M. (Jeroen))72–83 (Springer, Berlin, Heidelberg, 2007). doi:10.1007/978-3-540-75538-8_7.

41. Kocsis, L. & Szepesvári, C. Bandit Based Monte-Carlo Planning. in Machine Learning: ECML 2006 (eds. Fürnkranz, J., Scheffer, T. & Spiliopoulou, M.) 282–293 (Springer, Berlin, Heidelberg, 2006). doi:10.1007/11871842_29.

42. Vaswani, A. et al. Attention Is All You Need. Preprint at 10.48550/arXiv.1706.03762 (2023).

43. Glorot, X. & Bengio, Y. Understanding the difficulty of training deep feedforward neural networks. in Proceedings of the Thirteenth International Conference on Artificial Intelligence and Statistics 249–256 (JMLR Workshop and Conference Proceedings, 2010).

44. Rosenberg, S. A. et al. Identification of BING-4 Cancer Antigen Translated From an Alternative Open Reading Frame of a Gene in the Extended MHC Class II Region Using Lymphocytes From a Patient With a Durable Complete Regression Following Immunotherapy. The Journal of Immunology 168, 2402–2407 (2002).

45. Stewart-Jones, G. B. E., McMichael, A. J., Bell, J. I., Stuart, D. I. & Jones, E. Y. A structural basis for immunodominant human T cell receptor recognition. Nat Immunol 4, 657–663 (2003).

46. Cole, D. K. et al. Structural Mechanism Underpinning Cross-reactivity of a CD8+ T-cell Clone That Recognizes a Peptide Derived from Human Telomerase Reverse Transcriptase*. Journal of Biological Chemistry 292, 802–813 (2017).

47. Sachs, A. et al. Impact of Cysteine Residues on MHC Binding Predictions and Recognition by Tumor-Reactive T Cells. The Journal of Immunology 205, 539–549 (2020).

48. Holland, C. J. et al. Specificity of bispecific T cell receptors and antibodies targeting peptide-HLA. J Clin Invest 130, 2673–2688 (2020).

49. Sami, M. et al. Crystal structures of high affinity human T-cell receptors bound to peptide major histocompatibility complex reveal native diagonal binding geometry. Protein Engineering, Design and Selection 20, 397–403 (2007).

50. Knight, R. R. et al. A distinct immunogenic region of glutamic acid decarboxylase 65 is naturally processed and presented by human islet cells to cytotoxic CD8 T cells. Clinical and Experimental Immunology 179, 100–107 (2015).

51. Wu, D. et al. Structural assessment of HLA-A2-restricted SARS-CoV-2 spike epitopes recognized by public and private T-cell receptors. Nat Commun 13, 19 (2022).

52. Savage, S. R. et al. Pan-cancer proteogenomics expands the landscape of therapeutic targets. Cell 187, 4389-4407.e15 (2024).

53. Qudaihi, G. A. et al. Identification of a novel peptide derived from the M-phase phosphoprotein 11 (MPP11) leukemic antigen recognized by human CD8+ cytotoxic T lymphocytes. Hematology/Oncology and Stem Cell Therapy 3, 24 (2010).

54. Yang, F. et al. Validation of an IFN-gamma ELISpot assay to measure cellular immune responses against viral antigens in non-human primates. Gene Ther 29, 41–54 (2022).

55. Chersi, A., Modugno, F.di & Rosano, L. Flexibility of Amino Acid Residues at Position Four of Nonapeptides Enhances Their Binding to Human Leucocyte Antigen (HLA) Molecules. Zeitschrift für Naturforschung C 55, 109–114 (2000).

56. Vacic, V., Iakoucheva, L. M. & Radivojac, P. Two Sample Logo: a graphical representation of the differences between two sets of sequence alignments. Bioinformatics 22, 1536–1537 (2006).

