## Supplementary material for "Reengineering the antigen optimization process for superior neoantigen vaccine design": Cover letter

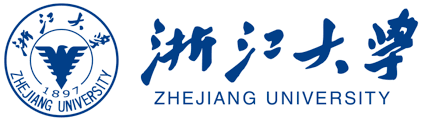
Qiushi Chair Professor and Director
Institute of Quantitative Biology
Zhejiang University, China

August 4^th^, 2025

Dear *BioRxiv* editors,

We are pleased to submit the manuscript titled “**Reengineering the antigen optimization process for superior neoantigen vaccine design**” for your kind consideration of publication in *BioRxiv.* This work presents AlphaVacc, a novel deep reinforcement learning framework that redefines how immunogenic peptides are generated for neoantigen-based cancer vaccines.

Neoantigen vaccines, derived from tumor-specific mutations, are a promising direction in cancer immunotherapy. However, identifying highly immunogenic peptides remains a major bottleneck. Existing predictive models often exhibit limited precision, and generative models that rely on these predictions tend to overfit to narrow regions of sequence space, compromising diversity and candidate quality.

To overcome these limitations, we developed AlphaVacc, which directly generates optimized peptide candidates using a Monte Carlo tree search (MCTS)-guided peptide hallucination process (Figure R1a). Instead of relying on scoring functions from external predictors, AlphaVacc autonomously learns how to modify peptides toward higher immunogenic potential. Central to this process is a supervised fine-tuning network (SFT), which guides mutation decisions within the MCTS loop (Figure R1b). The model is iteratively refined through reinforcement learning cycles, leveraging predictive feedback from prior optimization rounds and informed by IEDB peptide data (Figure R1c).

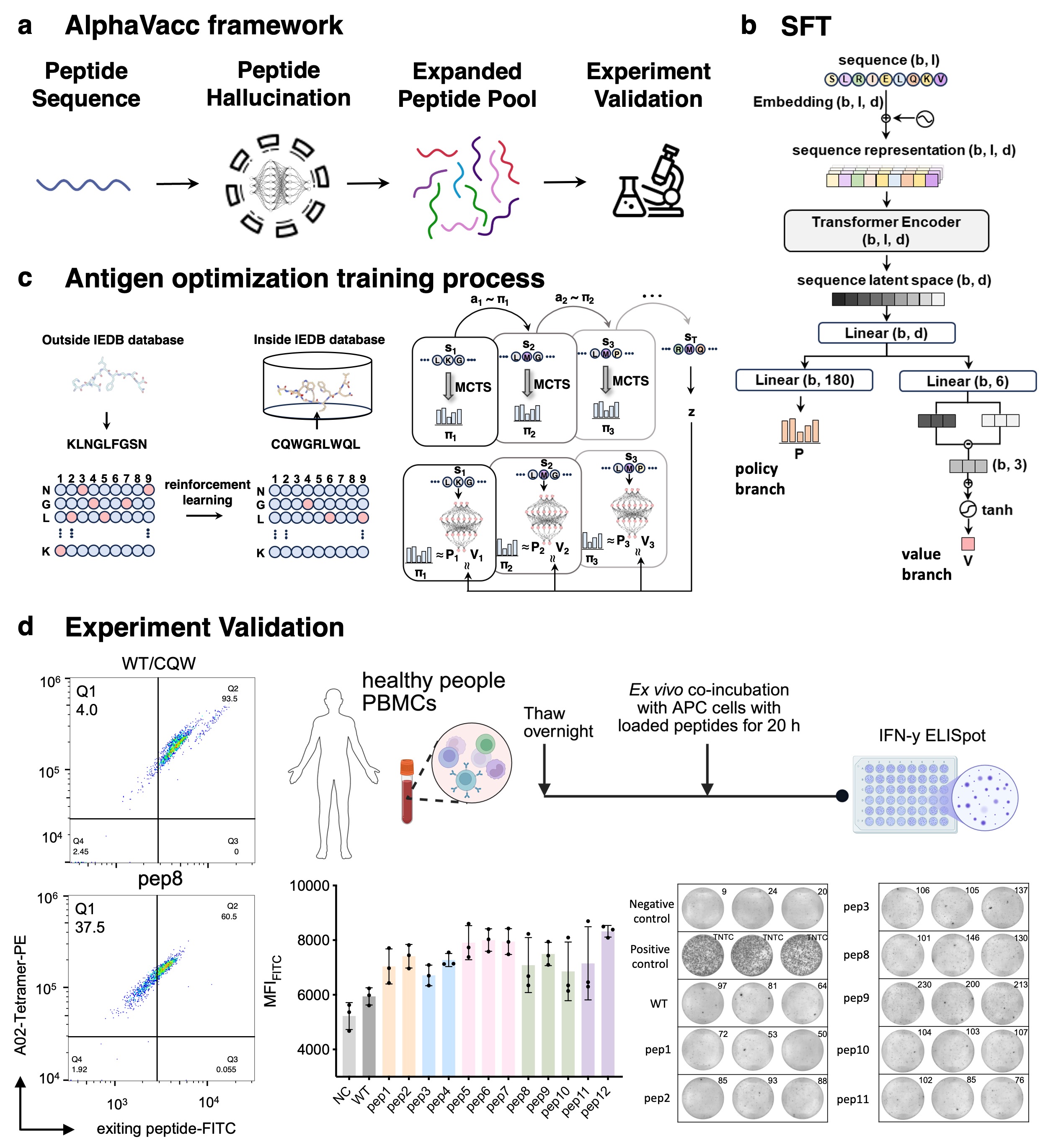

**Figure R1. Overview of the AlphaVacc framework and experimental validation. (a)** AlphaVacc optimizes peptides through an MCTS-guided peptide hallucination process, generating expanded peptide pool for experimental testing. **(b)** The structure of SFT(AlphaVacc) is illustrated. **(c)** AlphaVacc optimizes random peptides toward known immunogenic targets or until a step limit is reached; collected trajectories are used to retrain SFT(AlphaVacc). **(d)** Experimental validation of 12 AlphaVacc-generated peptides compared to the wild-type CQW peptide (WT).

Comprehensive in silico assessments demonstrated that AlphaVacc outperforms existing models in generating peptides with enhanced immunogenicity. In a case study, we optimized a BING-4 peptide CQWGRLWQL (CQW) to generate 12 candidate vaccines. Of these, 11 exhibited improved HLA-A*02:01 binding, as measured by flow cytometry, and 7 induced significant T-cell responses in IFN-γ ELISpot assays (Figure R1d). Detailed analysis revealed that AlphaVacc employs two distinct mutation strategies, *surge* and *progressive*, to efficiently guide peptides toward strong-binding states. Further experimental validation of 16 mutation groups confirmed 15 of AlphaVacc’s predictions, underscoring the model’s interpretability and reliability.

To conclude, our results demonstrate that AlphaVacc offers a robust and interpretable solution for peptide design, enabling efficient discovery of immunogenic candidates through learned mutation strategies. We believe this work will be of strong interest to the machine learning and biomedical communities, particularly in the context of vaccine development and computational immunology. It aligns with *BioRxiv*’s vision of advancing machine learning research that delivers meaningful impact across scientific disciplines.

Thank you for your time and consideration. We look forward to the opportunity to contribute to your journal.

Sincerely,

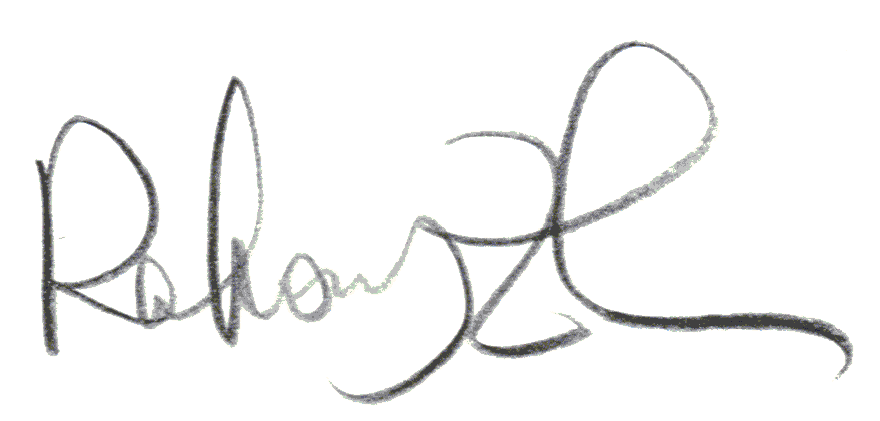

Ruhong Zhou, PhD

Zhejiang Univ and Columbia Univ
