## Supplementary Information-AlphaVacc-bioRxiv for "Reengineering the antigen optimization process for superior neoantigen vaccine design"

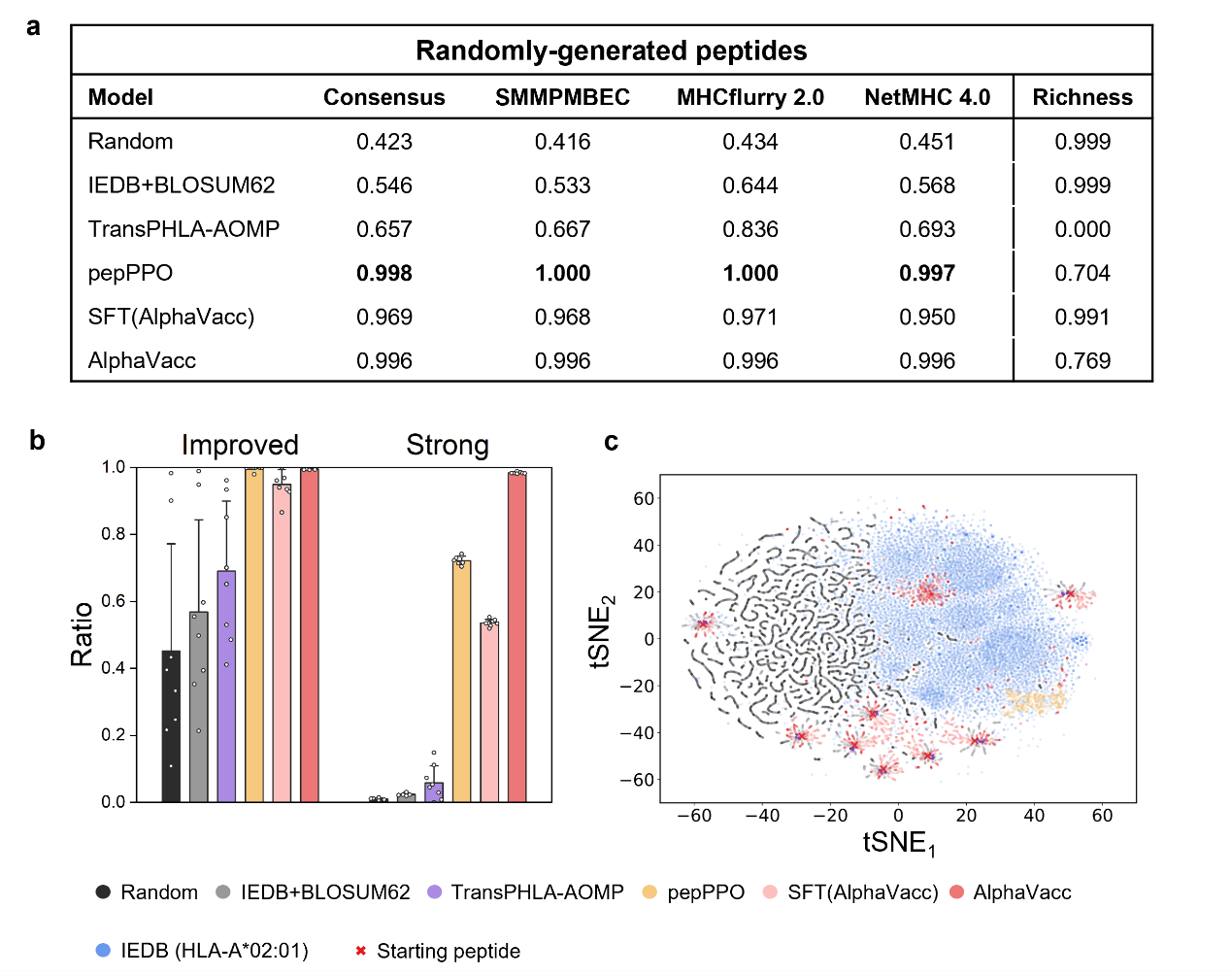

**Figure S1 Peptide optimization quality comparison between AlphaVacc and other baseline models using randomly-generated peptides. (a)** Different evaluation algorithms are used to assess the improved binder ratio in datasets generated by all the models. **(b)** A further comparison of the improved binder ratio and strong binder ratio among the different models. **(c)** The distribution of datasets generated by all the models, compared to that of the IEDB database.

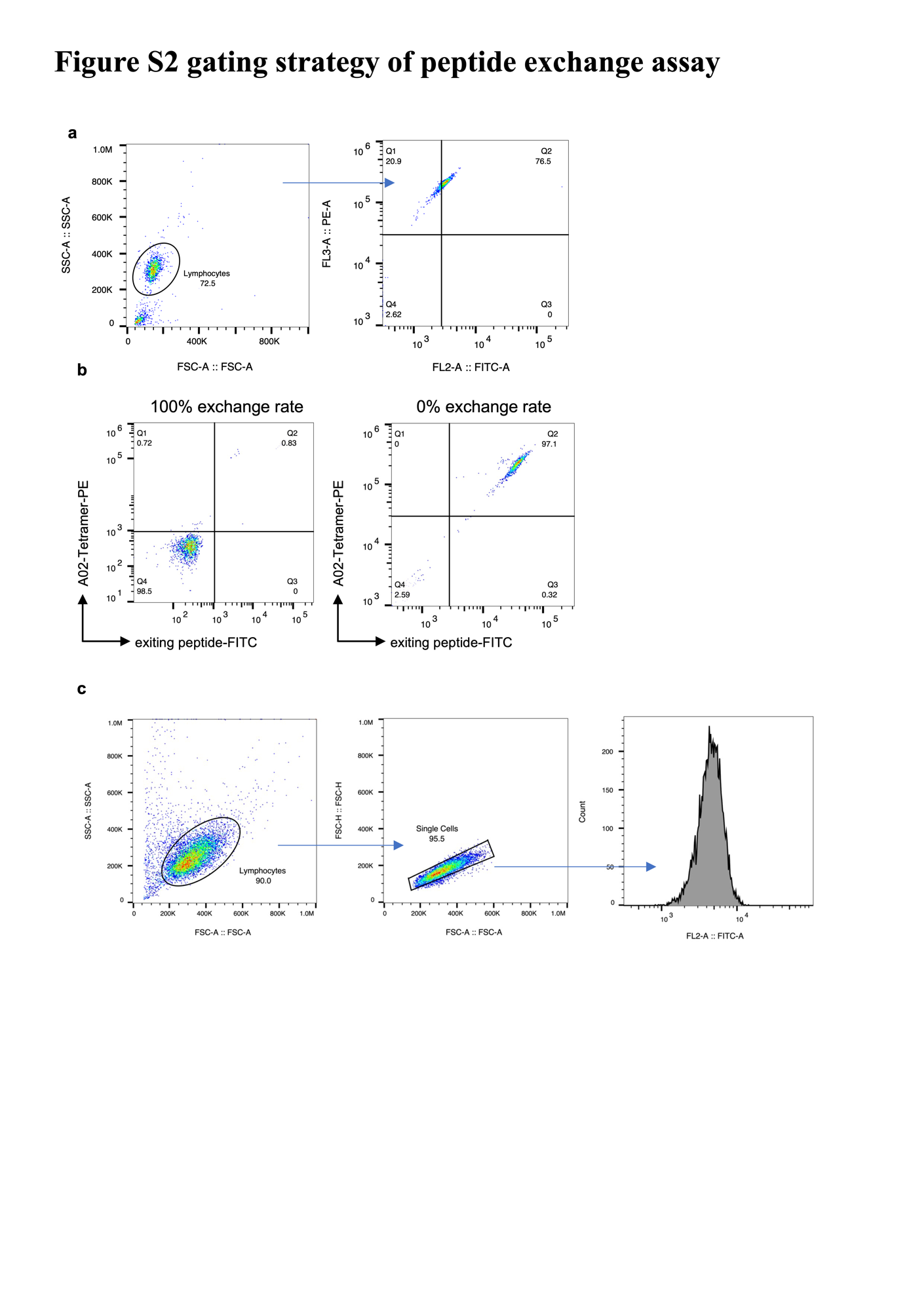

**Figure S2 Gating strategy for the CQW and candidate peptide exchange assay and T2 binding assay. (a)** Gating strategy for the peptide exchange assay. **(b)** Flow cytometric plots of 0% and 100% exchange rates, respectively. **(c)** Gating strategy of the T2 binding assay.

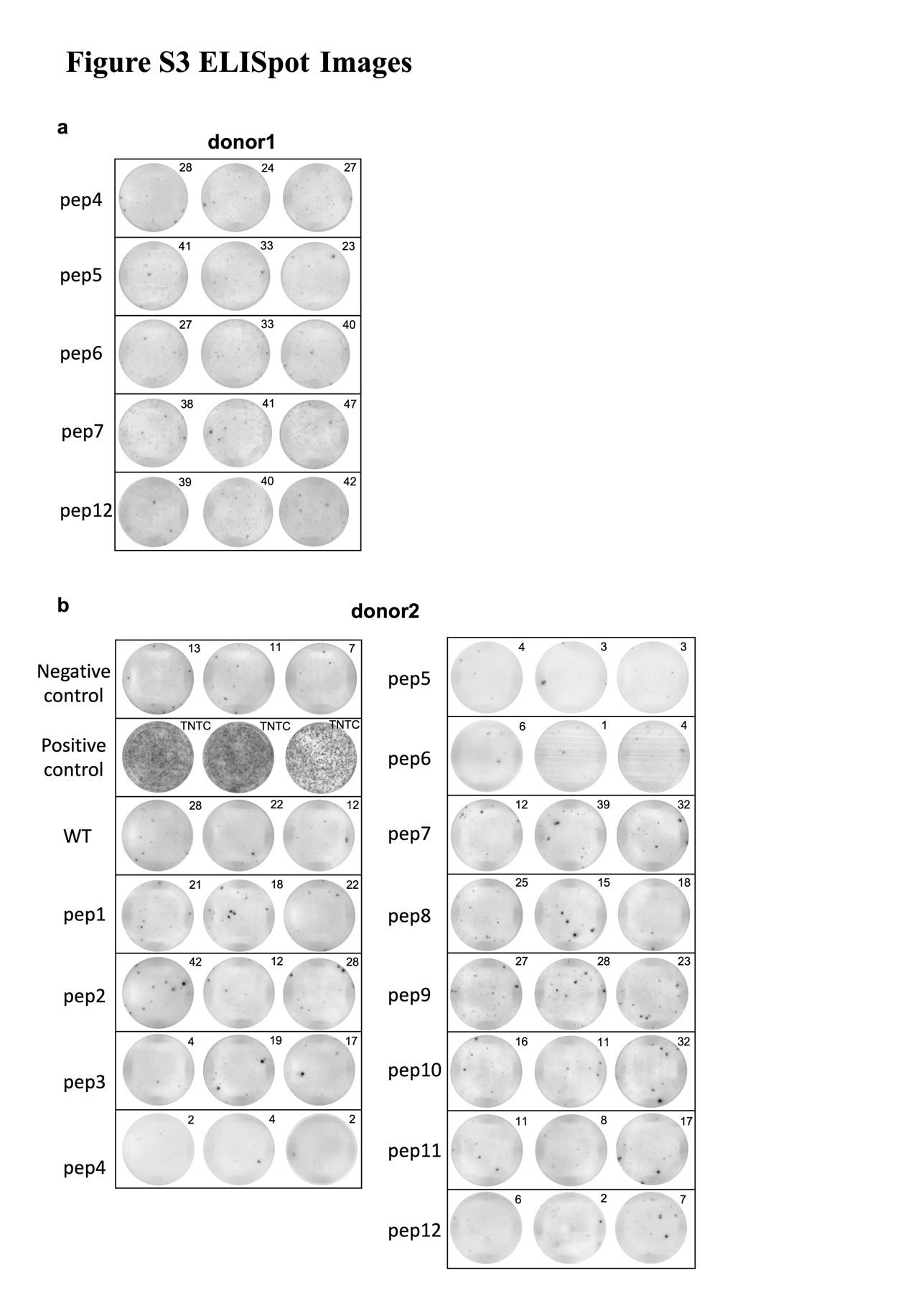

**Figure S3 ELISpot images. (a)** ELISpot images from donor 1 with no obvious activation. **(b)** ELISpot images from donor 2 show the same activation trend as donor 1.

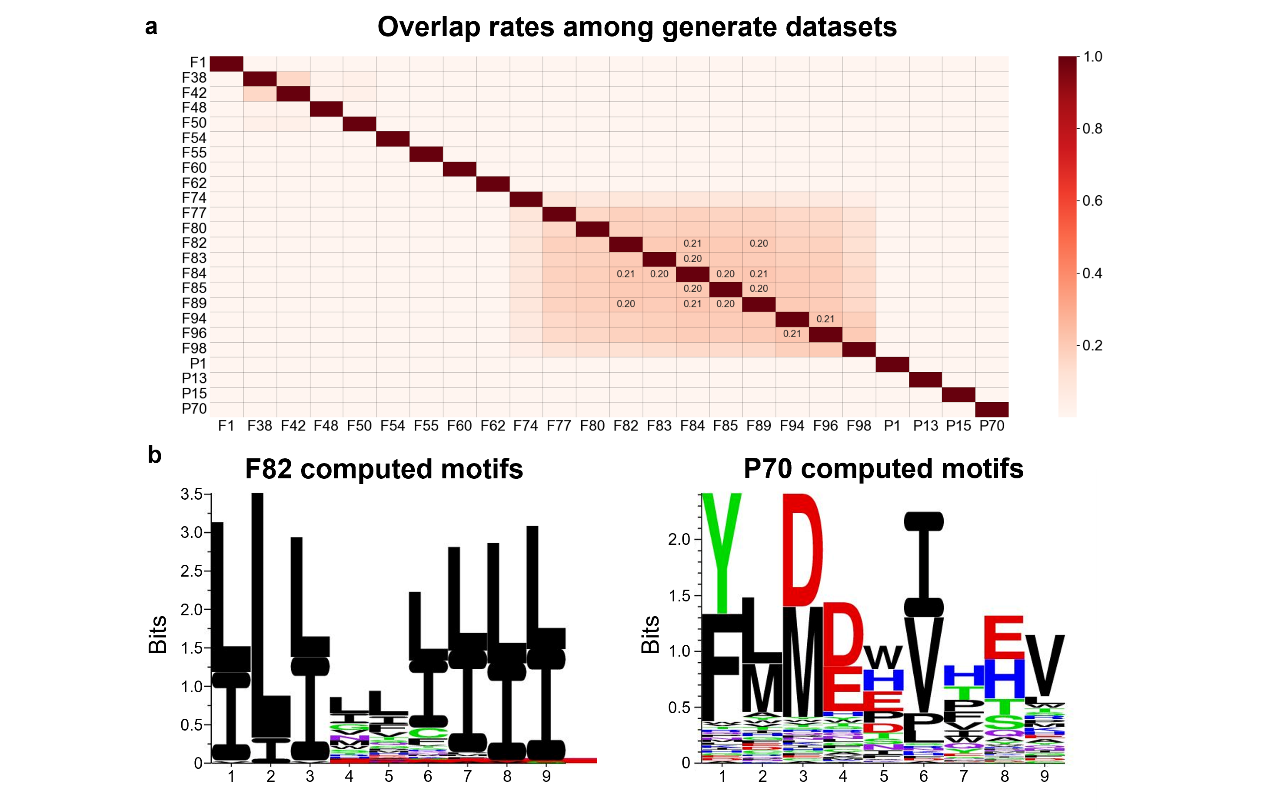

**Figure S4 Details related to the model winners of different rounds in arena competition process. (a)** The overlap rates among the generated datasets of different model winners. **(b)** The computed motifs of the generated datasets of models ‘F82’ and ‘P70’.

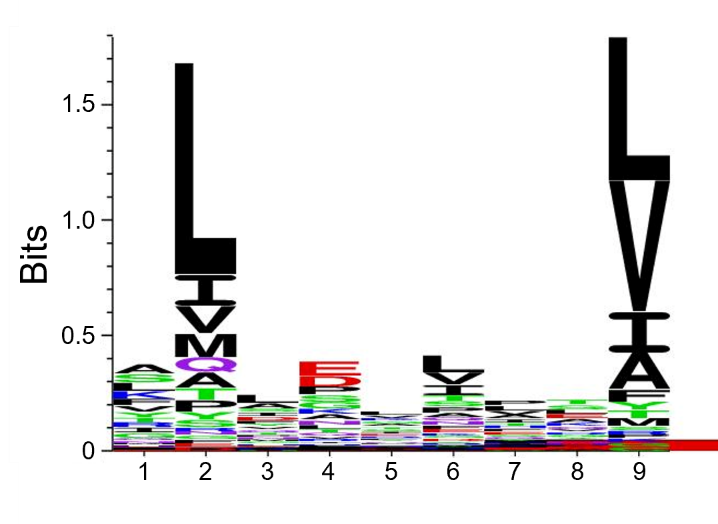

**Figure S5** The computed motifs of IEDB for HLA-A*02:01.

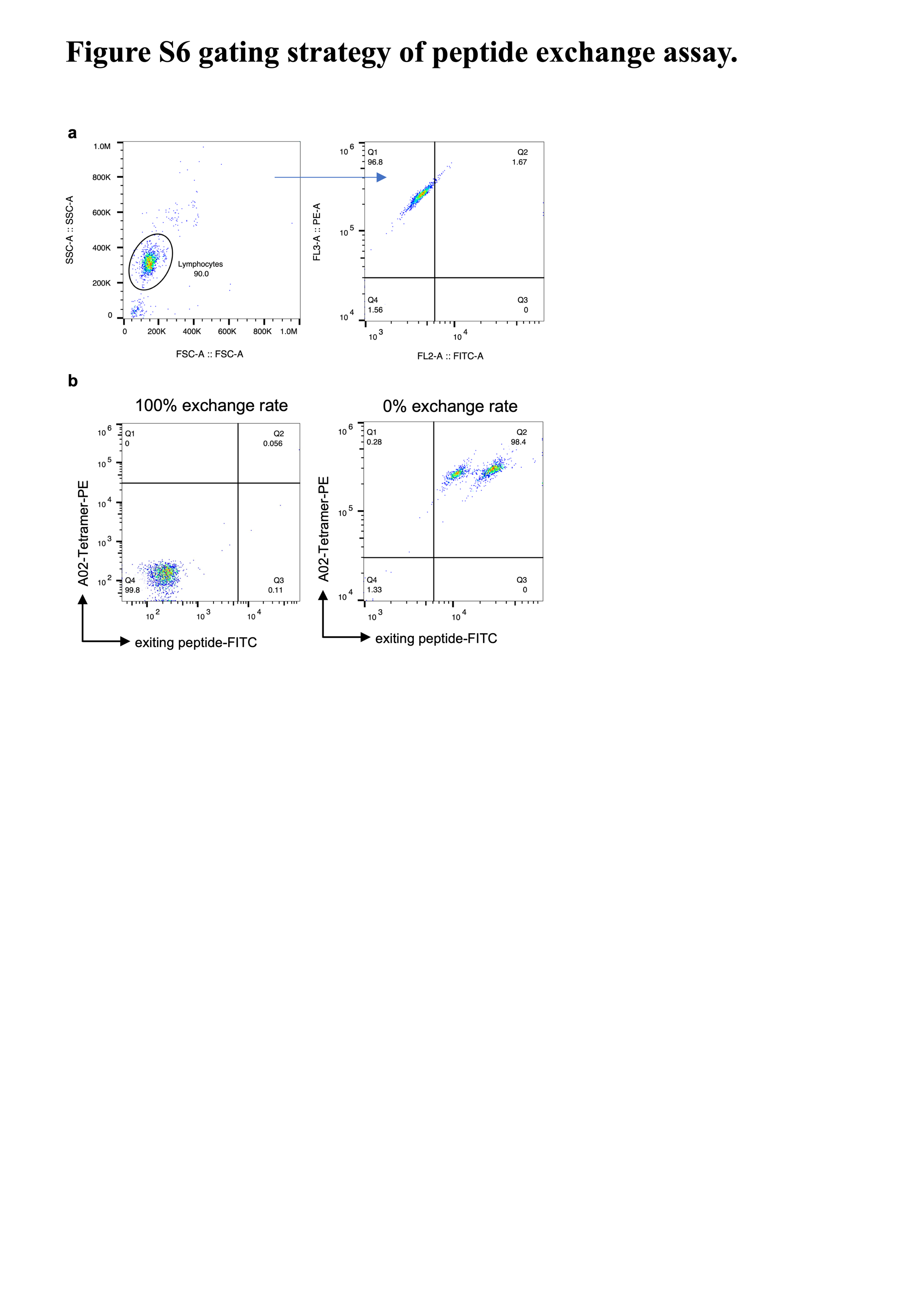

**Figure S6 Gating strategy of the peptide exchange assay in strategy validation. (a)** Gating strategy for Peptide exchange assay. **(b)** Flow cytometric plots of 0% and 100% exchange rates, respectively.

**Table S1 Peptide exchange efficiency of CQW and candidates.**

| **Analyzed sample** | **MFI_FITC_** |
| --- | --- |
| Control #2: 0% Exiting peptide or 100% peptide exchange | 449 |
| Control #3: 100% Exiting peptide or 0% peptide exchange | 35445 |

| **Analyzed sample** | **MFI_FITC_** | **%Peptide exchange efficiency** |
| --- | --- | --- |
| Standard peptide | 2903 | 92.99 |
| WT/CQW | 4634 | 88.04 |
|  | 4383 | 88.76 |
|  | 4903 | 87.27 |
| pep1 | 2777 | 93.34 |
|  | 3315 | 91.81 |
|  | 3359 | 91.68 |
| pep2 | 3945 | 90.01 |
|  | 4239 | 89.17 |
|  | 3913 | 90.10 |
| pep3 | 3137 | 92.32 |
|  | 3060 | 92.54 |
|  | 2980 | 92.77 |
| pep4 | 3173 | 92.22 |
|  | 3396 | 91.58 |
|  | 3511 | 91.25 |
| pep5 | 3520 | 91.22 |
|  | 3488 | 91.32 |
|  | 3223 | 92.07 |
| pep6 | 3449 | 91.43 |
|  | 3199 | 92.14 |
|  | 3581 | 91.05 |
| pep7 | 3706 | 90.69 |
|  | 3632 | 90.90 |
|  | 3779 | 90.48 |
| pep8 | 3297 | 91.86 |
|  | 2965 | 92.81 |
|  | 3137 | 92.32 |
| pep9 | 4486 | 86.32 |
|  | 5158 | 84.04 |
|  | 4851 | 85.08 |
| pep10 | 3274 | 90.44 |
|  | 3604 | 89.32 |
|  | 3850 | 88.49 |
| pep11 | 3083 | 91.09 |
|  | 2652 | 92.56 |
|  | 3816 | 88.60 |
| pep12 | 3504 | 89.66 |
|  | 3493 | 89.70 |
|  | 3650 | 89.17 |

**Table S2 MFI_FITC_ of all peptides in T2 binding assay.**

| **Analyzed sample** | **MFI_FITC_** | | |
| --- | --- | --- | --- |
| Only APC | 4677 | 5366 | 5628 |
| WT/CQW | 5610 | 6233 | 5982 |
| pep1 | 6432 | 6975 | 7711 |
| pep2 | 7823 | 7447 | 6962 |
| pep3 | 6700 | 6342 | 7090 |
| pep4 | 7137 | 7129 | 7550 |
| pep5 | 7386 | 8603 | 7736 |
| pep6 | 7580 | 8403 | 8030 |
| pep7 | 7484 | 8426 | 7938 |
| pep8 | 6346 | 8232 | 6684 |
| pep9 | 7098 | 7944 | 7437 |
| pep10 | 6140 | 8093 | 6344 |
| pep11 | 6298 | 8695 | 6462 |
| pep12 | 8499 | 8387 | 8090 |

**Table S3 Weak binder sequences and its corresponding strong binder sequences.**

| **Weak Binder**  **Sequence** |  | **Strong Binder**  **Sequence** |  |
| --- | --- | --- | --- |
| YMPKYVISE | Z1 | YMPKYVISV | Z2 |
| YVMPWIVLD | Z3 | YVMPWIVLV | Z4 |
| FLIYPVPTN | Z5 | FLIYPVPTV | Z6 |
| ALVEYVPTH | Z7 | ALVEYVPTV | Z8 |
| THMDLIILV | Z9 | TLMDLIILV | Z10 |
| YKIPALIAV | Z11 | YMIPALIAV | Z12 |
| FDMDHPTYL | Z13 | FLMDHPTYL | Z14 |
| FIMGHVLEW | Z15 | FIMGHVLEV | Z16 |

**Table S4 Normal binder sequences and its corresponding strong binder sequences.**

| **Normal Binder**  **Sequence** |  | **Strong Binder**  **Sequence** |  |
| --- | --- | --- | --- |
| ALRDILPHL | Z17 | ALMDILPHL | Z18 |
| KADDIRIFV | Z19 | KAMDIRIFV | Z20 |
| YIHDPVQHV | Z21 | YIMDPVQHV | Z22 |
| NLIGIRVQV | Z23 | NLIGIRVYV | Z24 |
| YLMGILIPE | Z25 | YLMDILIPE | Z26 |
| NLANPIVHV | Z27 | NLANPIFHV | Z28 |
| FAIDPVVHL | Z29 | FAIDLVVHL | Z30 |
| NLIFHQIIV | Z31 | NLIEHQIIV | Z32 |

**Table S5 Peptide exchange efficiency of all peptides.**

| **Analyzed sample** | **MFI_FITC_** |
| --- | --- |
| Control #2: 0% Exiting peptide or 100% peptide exchange | 395 |
| Control #3: 100% Exiting peptide or 0% peptide exchange | 22208 |

| **Analyzed sample** | **MFI_FITC_** | **%Peptide exchange efficiency** |
| --- | --- | --- |
| Standard peptide | 3535 | 85.60 |
| Z1 | 18886 | 15.23 |
| Z2 | 3727 | 84.72 |
| Z3 | 10371 | 54.27 |
| Z4 | 4269 | 82.24 |
| Z5 | 8357 | 63.5 |
| Z6 | 3630 | 85.17 |
| Z7 | 15329 | 31.54 |
| Z8 | 3594 | 85.33 |
| Z9 | 6360 | 72.65 |
| Z10 | 3335 | 86.52 |
| Z11 | 12649 | 43.82 |
| Z12 | 3260 | 86.87 |
| Z13 | 11739 | 47.99 |
| Z14 | 4521 | 81.08 |
| Z15 | 13392 | 40.42 |
| Z16 | 3630 | 85.17 |
| Z17 | 8028 | 65.01 |
| Z18 | 3096 | 87.62 |
| Z19 | 3465 | 85.93 |
| Z20 | 3228 | 87.01 |
| Z21 | 3890 | 83.98 |
| Z22 | 3399 | 86.23 |
| Z23 | 3741 | 84.66 |
| Z24 | 3917 | 83.85 |
| Z25 | 5188 | 78.03 |
| Z26 | 3385 | 86.29 |
| Z27 | 4685 | 80.33 |
| Z28 | 3870 | 84.07 |
| Z29 | 8056 | 64.88 |
| Z30 | 3708 | 84.81 |
| Z31 | 9461 | 58.44 |
| Z32 | 3630 | 85.17 |
